# Substrate induced activation in the conserved ribonuclease YicC

**DOI:** 10.64898/2026.05.21.726821

**Authors:** Kai Katsuya-Gaviria, Giulia Paris, Ben F. Luisi, Aleksei Lulla

## Abstract

YicC is a conserved protein, recently discovered to have hydrolytic endoribonuclease activity, with a phylogenetic distribution implicating an ancestry deeper than that for most extant ribonucleases central to RNA metabolism. We present evidence that *Escherichia coli* YicC can cleave the small regulatory RNAs (sRNAs), RyhB and RprA, even when they are sequestered in otherwise protective complexes with the RNA chaperone Hfq. Nonetheless, YicC activity is markedly diminished when RyhB and other sRNAs are paired with cognate mRNA. Our cryoEM structures of catalytically inactive YicC in complex with RyhB and in the *apo* state reveal quaternary and tertiary structural switches, triggered by substrate engagement, that engulf and ratchet a stem loop element of the sRNA into an internal channel, where metal-assisted hydrolytic action occurs. Based on these findings, we propose that the enzyme favours specific stem-loop structures and may discriminate between pools of active, target-engaged sRNAs and those that are inactive. The mechanism for substrate-triggered conformational switching could represent an ancient strategy for selective RNA degradation.

## Introduction

The gene encoding YicC was first reported as *orfY*, which, when deleted in tandem with the neighbouring *orfX* (*dinD*), lead to marked phenotypes in *Escherichia coli*: aberrant cell morphology during stationary phase and reduced viability at high temperatures (Poulsen & Jensen, 1991). For nearly three decades, YicC remained a protein of unknown function, until Chen *et al*. (2021) discovered that its overexpression reduced cellular abundance of small regulatory RNAs (sRNAs) (Chen *et al*., 2021). One key sRNA affected was RyhB, a posttranscriptional regulator of iron stress responses, and overexpression of YicC was accompanied by a corresponding boost in expression of those genes otherwise suppressed by RyhB. Notably, RyhB abundance did not significantly change upon YicC overexpression in a strain lacking the exoribonuclease polynucleotide phosphorylase (PNPase), suggesting that YicC functions not as a nuclease itself, but as an adaptor, perhaps guiding PNPase to degrade RyhB (Chen *et al*., 2021). However, subsequent studies have demonstrated ribonuclease activity for purified *E. coli* YicC and for a *Bacillus subtilis* homologue, YloC (Huang *et al*., 2023; Ingle *et al*., 2022; Wu *et al*., 2024). In *Clostridioides difficile*, the sRNA SQ528 is a putative sRNA target of a YicC-like protein, underscoring a potential role of YicC and its homologues in sRNA regulation across diverse bacterial species (Martins *et al*., 2025). These and other observations raise the possibility that YicC might help to control levels of RyhB and other sRNAs.

The primary biological role of RyhB, in bacteria such as *E. coli* and closely related species, is to coordinate a system-wide post-transcriptional downregulation of non-essential iron-using proteins (*e.g.* SodB, SdhC) and upregulation of iron uptake pathways (*e.g.* ShiA, CirA) (Chareyre & Mandin, 2018; Prévost *et al*., 2007; Salvail *et al*., 2013). By controlling the expression of over 140 genes, RyhB helps to maintain core metabolic functions and survival during prolonged periods of iron scarcity. This regulation is critical for bacterial fitness and pathogenesis, as iron acquisition is a key competition between microbes and their hosts (Chareyre & Mandin, 2018). Like many other *trans*-acting sRNAs, RyhB actions rely on the assistance of RNA chaperones like Hfq, which acts as a cellular reservoir for sRNAs, binding them and providing protection from indiscriminate degradation (Vogel & Luisi, 2011; Roca *et al*., 2025; Santiago-Frangos & Woodson, 2018). By sequestering sRNA-mRNA complexes, Hfq can either sterically hinder access to cleavage sites, thereby protecting the RNAs, or conversely, it can promote conformational changes that present cleavage sites to an incoming ribonuclease, thereby actively promoting degradation (Bandyra *et al*., 2024; Dendooven *et al*., 2021).

Here, we characterise the direct endoribonucleolytic cleavage of RyhB by YicC and the impact of Hfq to guide its cleavage. Our cryoEM structures of the ribonuclease in its *apo* state reveal a dynamic "breathing" equilibrium that maintains the ribonuclease in an inactive surveillance mode, while structures of the RyhB-bound complex shed light on the structural basis for sRNA recognition and engagement. We also show that YicC directly cleaves the sRNA RprA *in vitro*, demonstrating its capacity to process additional sRNA targets. Furthermore, we show that sRNA–mRNA base-pairing inhibits YicC cleavage, potentially allowing YicC to discriminate between active, target-engaged sRNAs, and inactive sRNA pools.

## Results

### Conserved sequence patterns in YicC proteins

YicC-like proteins have been identified in diverse bacterial phyla, as represented by species such as Gram-negative *E. coli* and Gram-positive *B. subtilis* and *C. difficile* (Chen *et al*., 2021; Huang *et al*., 2023; Ingle *et al*., 2022; Martins *et al*., 2025; Wu *et al*., 2024). Through a comprehensive genomic survey of the Genome Taxonomy Database (GTDB R214) we observed that the high degree of conservation across the bacterial kingdom extends well beyond these specific species and their close relatives (Appendix Fig. S1). Across 81 bacterial phyla with at least 10 genomes annotated, YicC homologues were present in 71 phyla (88%). Among the extensively sampled phyla (>1000 genomes annotated in the database), YicC-like proteins are prevalent across Pseudomonadota (71%), Bacillota (70%), Bacteroidota (87%) and Desulfobacterota (89%), but absent in annotated genomes in Cyanobacteriota, Campylobacterota. Sequence conservation patterns from the family highlight residues of potential function importance (Appendix Fig. S2), to which we will return when discussing our structural findings below. RyhB is not co-conserved with YicC, which suggests that the protein likely evolved other roles in RNA metabolism.

### E. coli YicC exhibits intrinsic endonucleolytic activity

Recent studies have demonstrated that YicC homologues from *E. coli* and *B. subtilis* possess intrinsic endonucleolytic activity against short (<30 nts) synthetic single-stranded RNA oligonucleotides (Chen *et al*., 2021; Huang *et al*., 2023; Ingle *et al*., 2022; Wu *et al*., 2024). Furthermore, the *C. difficile* YicC homologue (CD25890) has been shown to form a catalytically active complex with PNPase and to be able to cleave SQ528, a regulatory sRNA involved in sporulation (Martins *et al*., 2025). To test whether *E. coli* YicC might act solely as a PNPase adaptor or could independently cleave a full-length, structured sRNA, we performed *in vitro* degradation assays, incubating *in vitro*-transcribed RyhB with purified YicC, PNPase, or both, and quenching the reactions at various time points (Fig. 1A). Using an enzyme-to-substrate ratio of 1:1, PNPase completely degraded naked RyhB under phosphorolytic conditions, as expected. Notably, YicC was also capable of degrading RyhB in the absence of PNPase, and the two enzymes do not appear to act synergistically. Similar outcomes, albeit with slower kinetics, were observed at a 1:10 enzyme-to-substrate ratio (Appendix Fig. S3). The degradation of RyhB by YicC did not generate detectable nucleoside mono- or diphosphates, suggesting that the enzyme does not have hydrolytic or phosphorolytic exonuclease activity (Appendix Fig. S4).

**Figure 1:**
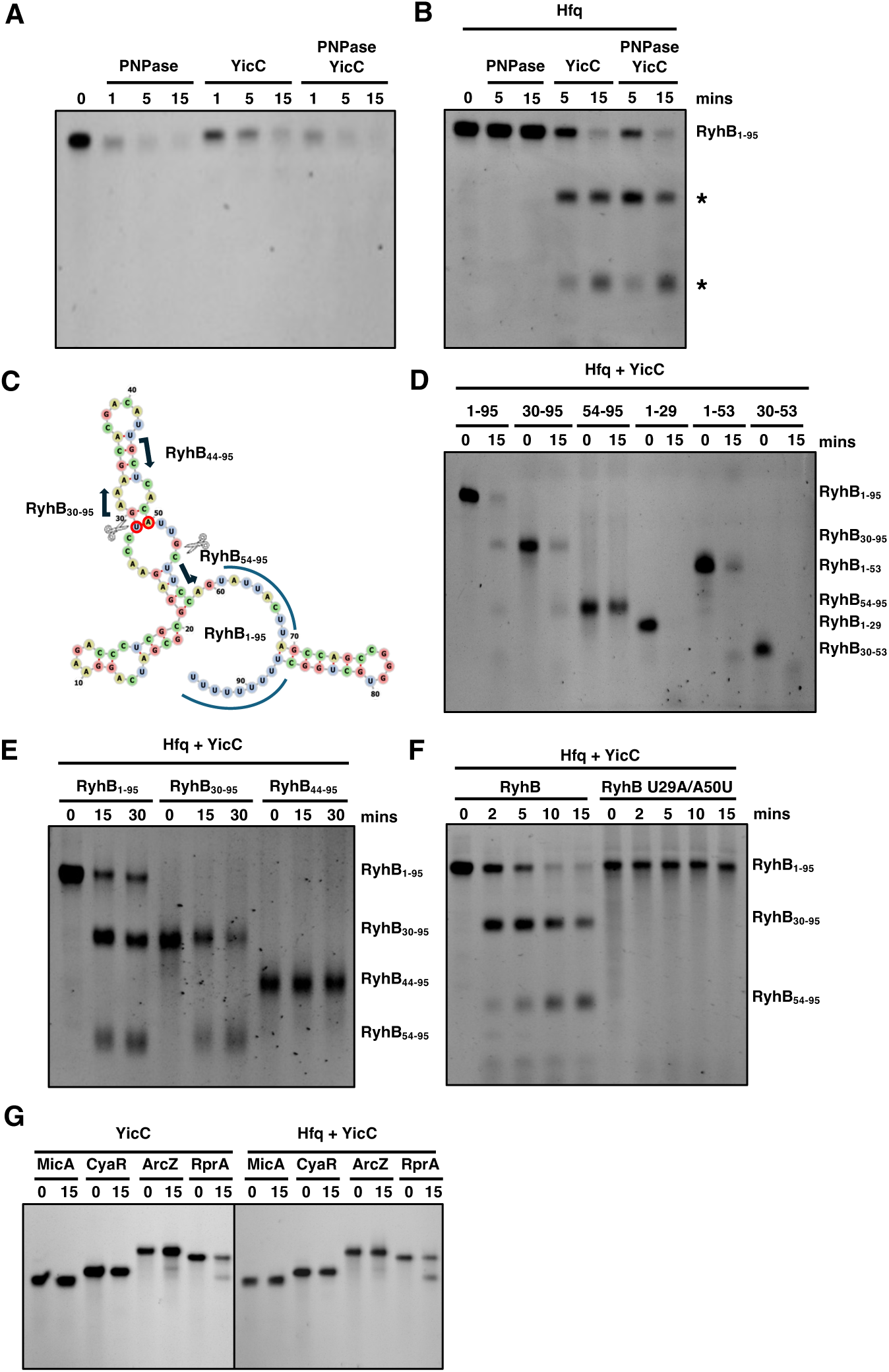
YicC degrades RyhB with substrate selectivity. *In vitro* degradation assays showing the stability of RyhB sRNA over a 15-minute time course. **(A,B)** RyhB was incubated with PNPase, YicC, or both proteins in the absence (A) and presence (B) of the RNA chaperone Hfq. While Hfq protects RyhB from PNPase, YicC can still cleave the sRNA, resulting in two stable cleavage products (marked with asterisks). **(C)** Secondary structure of RyhB (predicted with RNAfold, Gruber et al., 2008) showing the location of YicC cleavage sites (indicated by scissors icons). Blue lines indicate the nucleotides that interact with Hfq. Red circles indicate the nucleotides mutated in the U29A/A50U transversion. **(D)** *In vitro* degradation assays testing the susceptibility of the individual RyhB fragments after cleavage at sites identified by 5’ RACE. (Note: To allow for efficient T7 *in vitro* transcription, a single guanine was added to the 5’ end of the RyhB_54-95_ construct). **(E)** *In vitro* degradation assays evaluating structural requirements for YicC activity, showing that the RyhB_44-95_ fragment (lacking the upstream arm of the central stem) is resistant to cleavage. **(F)** *In vitro* degradation assays showing that the U29A/A50U mutation impairs YicC cleavage of Hfq-bound RyhB. **(G)** *In vitro* degradation assays testing the specificity of YicC against other sRNAs (MicA, CyaR, ArcZ, RprA) in the absence (left) and presence (right) of Hfq. YicC shows specificity, cleaving RprA but not the other sRNAs tested under these conditions.

### YicC bypasses Hfq-mediated protection

We next investigated the susceptibility of RyhB to YicC cleavage in the presence of the RNA chaperone Hfq. *In vivo*, Hfq facilitates RyhB-mediated regulation and generally protects this otherwise unstable sRNA from major cellular ribonucleases, including RNase E and PNPase (Vogel & Luisi, 2011). While YicC completely degrades naked RyhB, the presence of Hfq appears to restrict cleavage to specific sites, yielding two distinct, stable RNA fragments, which are potentially protected by the RNA chaperone (Fig. 1B). Hfq interacts with RyhB primarily via its 3’ poly-U tail (nucleotides 87-95) and makes additional contacts with an upstream UA-rich region (nucleotides 61-65), characteristic of Class I sRNAs (Fig. 1C). We hypothesised that this stable interaction preferentially protects the 3’ end of RyhB, that remains bound to Hfq post-cleavage, leaving the Hfq-free 5’-end product exposed to YicC cleavage.

Previous reports have shown that PNPase does not degrade RyhB when the sRNA is bound to Hfq; instead, it engages in a protective assembly with the Hfq-RyhB complex that precludes cleavage by alternative ribonucleases such as RNase E (Bandyra *et al*, 2016; Dendooven *et al*, 2021). Strikingly, YicC bypassed this Hfq- and Hfq-PNPase-mediated protection, cleaving the sRNA to generate the two stable, discrete products seen in the absence of PNPase (Fig. 1B). Taken together, these results indicate that YicC possesses intrinsic, PNPase-independent endonucleolytic activity against a full-length sRNA, and has the capacity to overcome Hfq and dual PNPase/Hfq protection, distinguishing it from other major cellular ribonucleases.

### YicC cleavage sites on RyhB, structural requirements and substrate specificity

To experimentally identify the precise cleavage sites of YicC on RyhB, we analysed the cleavage products by 5’ Rapid Amplification of cDNA Ends (5’ RACE), which identified sequences corresponding to cleavage immediately downstream of U29 or G53. To confirm these mapping results, we generated *in vitro*-transcribed RyhB fragments corresponding to the predicted cleavage products (RyhB_30-95_ and RyhB_54-95_) and tested their sensitivity to YicC in the presence of Hfq. As shown in Fig. 1D, these fragments recapitulated the sizes of the products generated from full-length RyhB (RyhB_1-95_). Notably, the RyhB_30-95_ fragment remained susceptible to the secondary cleavage event at G53, whereas the RyhB_54-95_ fragment was protected from further YicC activity by Hfq. *In vitro* transcribed RNA fragments corresponding to 5’ end products resulting from cleavage at these sites (i.e. RyhB_1-29,_ RyhB_1-53_, RyhB_30-53_) were also degraded by YicC (Fig. 1D).

Both identified cleavage sites (U29 and G53) reside within the predicted central stem-loop of RyhB (Fig. 1C). To determine whether this secondary structure is required for recognition, we synthesised RyhB_44-95_, a fragment containing the G53 cleavage site but lacking the upstream arm of the central stem. Interestingly, Hfq-bound RyhB_44-95_ was resistant to YicC cleavage (Fig. 1E). This indicates that, beyond primary sequence recognition, YicC likely requires a specific three-dimensional fold for substrate recognition and cleavage. To validate the U29 cleavage site within the context of the full-length sRNA, we introduced a U29A substitution, coupled with a compensatory A50U mutation to preserve the predicted secondary structure of the central stem-loop. The Hfq-bound RyhB U29A/A50U mutant was resistant to YicC cleavage compared to the wild-type sRNA (Fig. 1F).

Finally, we explored the broader substrate preferences of YicC. Chen *et al*. (2021) reported that YicC overexpression in *E. coli* reduces cellular levels of both RyhB and MicA (Chen *et al*, 2021). To determine whether this reduction is due to direct turnover or indirect regulatory effects, we incubated MicA with YicC *in vitro*. Unlike RyhB, MicA was not cleaved by YicC under our assay conditions (Fig. 1G). Furthermore, other Class I (ArcZ) and Class II (CyaR) sRNAs also proved resistant. RprA—an sRNA categorised as an intermediate between Class I and Class II—was, however, cleaved by YicC. In contrast to the effect observed on RyhB cleavage, the addition of Hfq did not alter the YicC cleavage pattern for RprA. These findings suggest that YicC acts with substrate selectivity, targeting specific sRNAs and structural motifs rather than acting as a general, promiscuous ribonuclease.

### Base-pairing protects sRNAs from YicC-mediated cleavage

Having established that YicC efficiently cleaves RyhB and RprA *in vitro*, we next investigated whether this nuclease activity is sensitive to the functional state of the sRNA, specifically, whether RNA-RNA base-pairing alters its susceptibility to YicC. We selected two distinct regulatory scenarios: the sequestration of RyhB by the RNA sponge ETS^LeuZ^, and the canonical base-pairing of RprA to the 5′ UTR of its target mRNA, *csgD* (Andreassen *et al*., 2018; Lalaouna *et al*., 2015).

The addition of the ETS^LeuZ^ sponge, which locks RyhB into an extended RNA-RNA duplex, almost completely abolished YicC activity over the same time course (Figs. 2A,B). This behaviour contrasts sharply with that of the canonical endoribonuclease RNase E. When we incubated RyhB and ETS^LeuZ^ with the catalytic core of RNase E (RNase E (1-598)), duplex formation actively promoted RyhB degradation, while the sRNA remained largely protected when bound to Hfq in the absence of the sponge (Figs. 2C,D).

**Figure 2.**
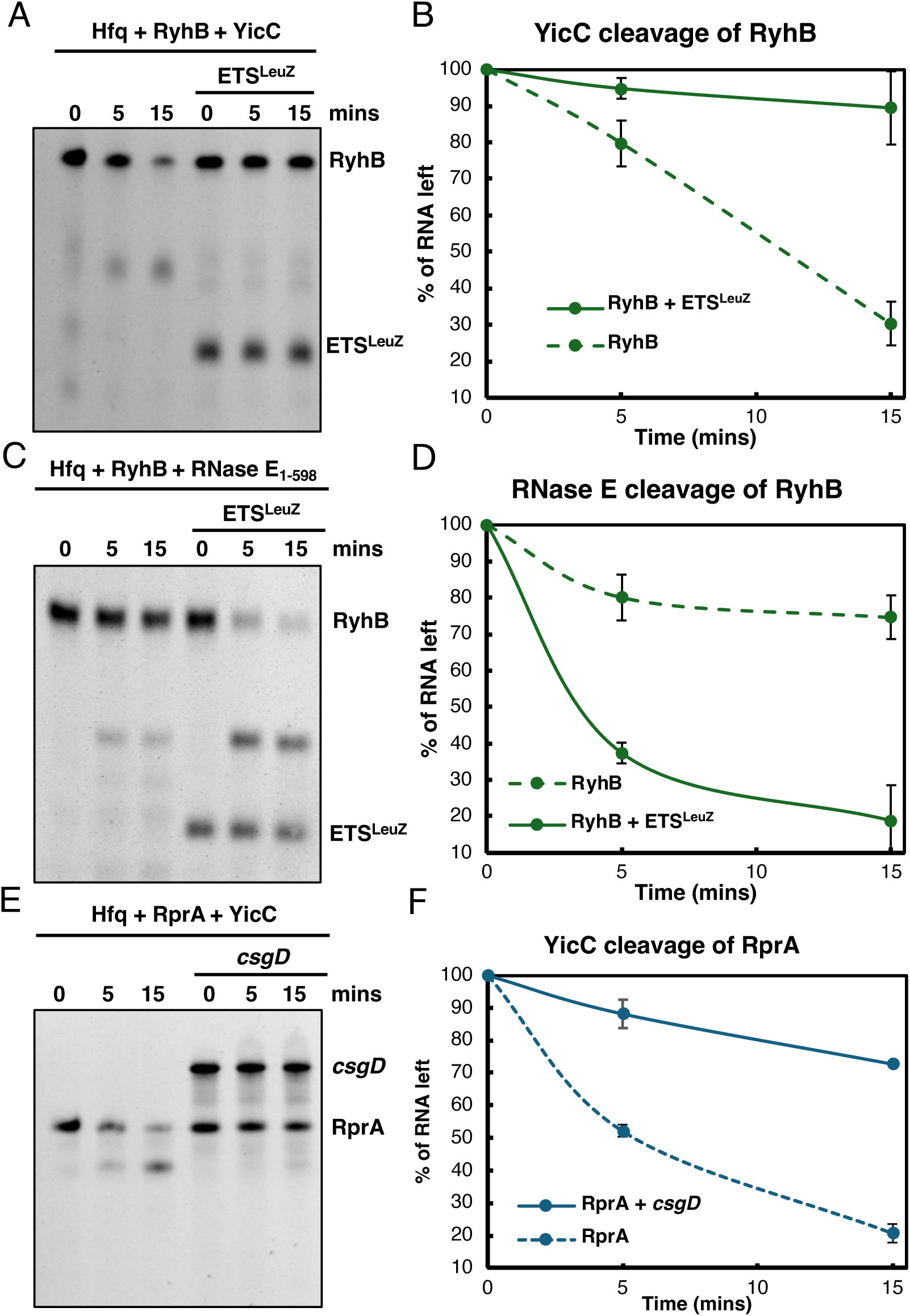
RNA-RNA base-pairing protects sRNAs from YicC endonucleolytic cleavage. (A,C) Representative *in vitro* cleavage assay showing the degradation of full-length RyhB by YicC (A) or RNase E (1-598) (C) in the presence of the RNA chaperone Hfq. The addition of the RNA sponge ETS^LeuZ^, which forms a stable duplex with RyhB, protects the sRNA from YicC cleavage over a 15-minute time course, while it promotes efficient cleavage by RNase E (1-598). (B,D) Quantification of RyhB degradation from the assays in A and C, respectively. The percentage of remaining full-length RNA is plotted against time for unengaged RyhB (dashed green line) and the RyhB-ETS^LeuZ^ duplex (solid green line). (E) Representative *in vitro* cleavage assay demonstrating the degradation of the sRNA RprA by YicC in the presence of Hfq. Canonical base-pairing with a fragment of its target mRNA, *csgD* (-147 to +18 relative to the translation start site), protects RprA from cleavage. (F) Quantification of RprA degradation from the assays in (E). The percentage of remaining full-length RNA is plotted against time for free RprA (dashed blue line) and the RprA-*csgD* complex (solid blue line). For both graphs, data points represent the mean ± standard deviation of n=3 independent replicates.

The protective effect against YicC cleavage was equally present in a canonical sRNA-mRNA silencing complex. Like RyhB, YicC efficiently cleaved free RprA, whereas the presence of a *csgD* mRNA fragment provided significant protection from degradation (Figs. 2E,F). Together, these data indicate that YicC preferentially targets unpaired sRNAs. The formation of an RNA duplex likely occludes the specific structural motifs required for YicC recognition or creates steric clashes that prevent the RNA from being accommodated at the active site. This suggests YicC may act as a clearance factor that selectively degrades excess or orphaned sRNAs without prematurely dismantling active regulatory complexes.

### YicC silences RyhB activity in vivo

Having observed that YicC efficiently cleaves RyhB *in vitro*, we next explored YicC action on RyhB *in vivo*. We adapted a three-component fluorescent system, in which the fluorescent protein mCherry is translationally fused to *sodB*, a well-characterised RyhB mRNA target (Chen *et al*., 2021). In the first component, the *sodB-mCherry* fusion is constitutively expressed, rendering the cells fluorescent red. In the second component, co-expression of RyhB represses the reporter, significantly reducing cellular fluorescence. The third component introduces YicC: if expressed and capable of inactivating RyhB, the ribonuclease relieves the downregulation of the *sodB-mCherry* fusion, thereby restoring the fluorescence signal.

RyhB-mediated repression of *sodB* occurs at two distinct levels: the sRNA binds at the ribosome binding site (RBS) to block translation initiation, and it subsequently induces cleavage of the *sodB* transcript by the RNA degradosome at a distal site approximately 350 nucleotides into the open reading frame (ORF) (Prevost *et al*., 2011). To capture this native regulatory mechanism, we designed our reporter to include the complete *sodB* ORF upstream of the *mCherry* fusion, enabling the distal cleavage event.

We first tested whether YicC could functionally clear RyhB in an *E. coli* wild-type background. The full-length *sodB-mCherry* construct, along with RyhB and YicC, were expressed from compatible plasmids under the control of the constitutive cp26 promoter. Strong constitutive expression of wild-type RyhB resulted in a growth defect (OD_600_ reduced to ∼1.25 compared to 1.87) and significantly repressed *sodB-mCherry* translation, confirming active target engagement (Fig. 3A). Co-expression of YicC increased reporter fluorescence and partially restored growth, consistent with *in vivo* inhibition of RyhB activity. We noted that high-level overexpression of YicC alone also reduced the basal fluorescence of the reporter in the empty-vector control strains, likely due to pleiotropic effects or non-specific transcript turnover associated with ribonuclease overexpression.

**Figure 3.**
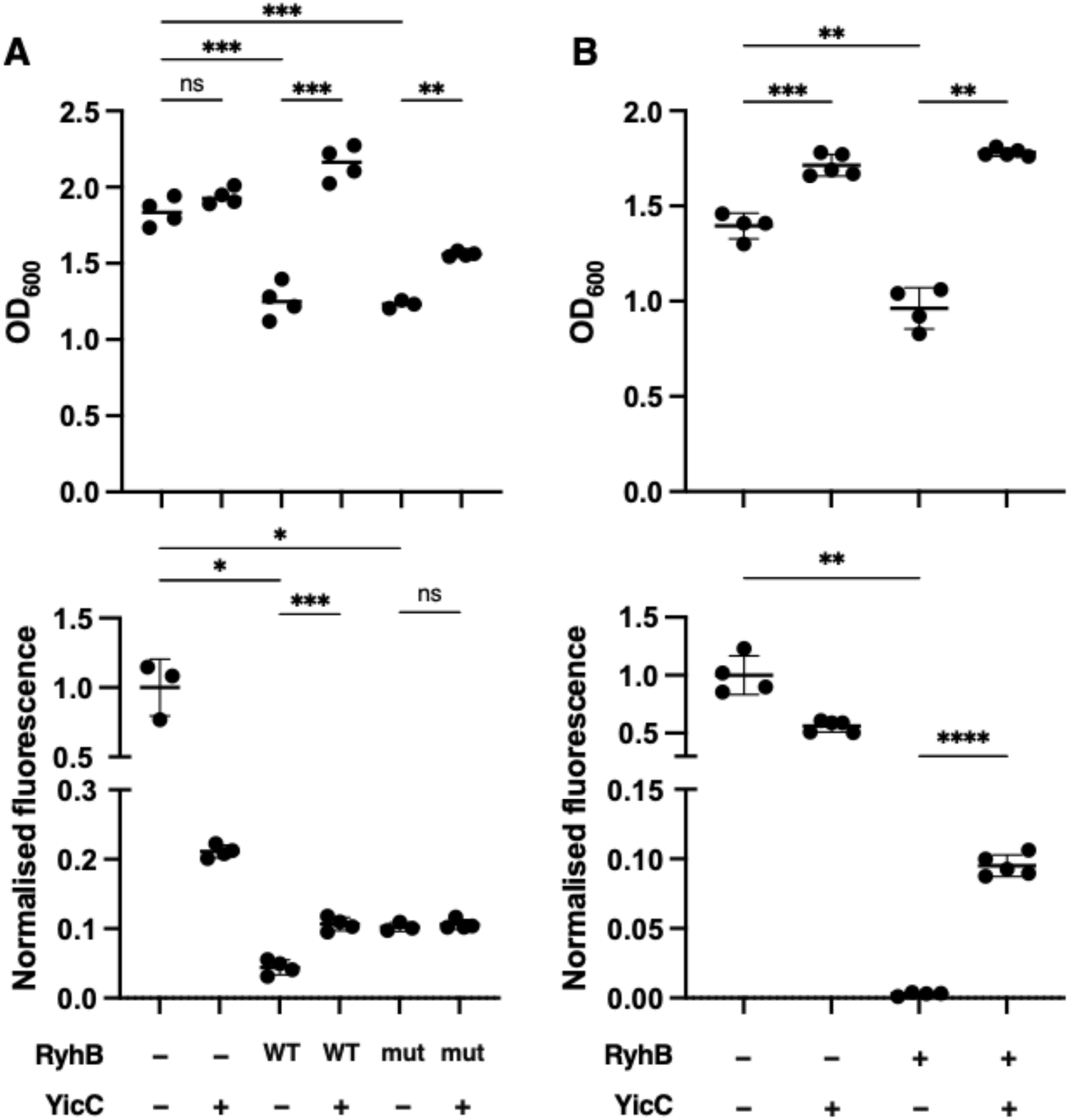
YicC inactivates RyhB *in vivo* independently of the exo-nuclease PNPase. (A) Growth (OD_600_, top) and normalised fluorescence (bottom) of the *sodB-mCherry* translational reporter in a wild-type *E. coli* background. Cells were co-transformed with compatible plasmids expressing either wild-type (WT) RyhB or the cleavage-resistant U29A/A50U mutant (mut), alongside either YicC (+) or an empty vector control (-). YicC co-expression significantly rescues both the growth defect and fluorescence repression caused by WT RyhB but fails to rescue fluorescence against the mutant sRNA. (B) Growth (OD_600_, top) and normalised fluorescence (bottom) in a Δ*pnp*Δ*rna* double deletion background. YicC maintains the capacity to robustly de-repress the *sodB-mCherry* reporter in the absence of PNPase and RNase I ribonucleases. For all graphs, data points represent individual biological replicates (n=4). Horizontal bars indicate the mean, and error bars represent the standard deviation. Statistical significance was determined using Welch’s ANOVA (not assuming equal variance) followed by Dunnett’s T3 multiple comparisons test (* p<0.05; ** p<0.01; *** p<0.001; **** p<0.0001; ns, not significant).

To test whether this rescue was structurally specific and not an artifact of YicC overexpression, we tested the RyhB U29A/A50U mutant, which we previously identified as resistant to YicC cleavage *in vitro*. The mutant sRNA retained biological activity, repressing the *sodB* reporter and restricting growth to similar levels as the wild-type sRNA. Notably, while co-expression of YicC provided a partial restoration of growth to these cells, it failed to rescue the reporter fluorescence showing that YicC cannot functionally clear the mutant sRNA, consistent with our *in vitro* results (Fig. 3A). Together, these data are consistent with the *in vitro* cleavage activity observed for YicC against full-length RyhB.

Finally, to test if the activity is PNPase-dependent, we used our reporter system in a double deletion strain lacking both PNPase and the non-specific endonuclease RNase I (Δ*pnp* Δ*rna*). Because RyhB baseline stability is altered in the absence of PNPase (Bandyra *et al*., 2016), expressing the sRNA from a constitutive promoter ensured sufficient steady-state levels to assess YicC’s impact. Notably, YicC remained able to de-repress the *sodB-mCherry* reporter in the Δ*pnp* Δ*rna* background, consistent with direct RyhB inactivation *in vivo* (Fig. 3B).

### A dynamic trimer-trimer interface stabilises apo-YicC conformations

Crystallographic analyses have revealed that *E. coli* YicC assembles into a hexamer in which two trimeric units swivel relative to each other through a scissoring rotation (Huang *et al*., 2023; Wu *et al*., 2024). Each protomer can be structurally decomposed into four principal domains, to which we will refer throughout as α/β, cap, four-helix bundle, and helical hairpin (Fig. 4A). Our cryoEM data corroborate these earlier findings and reveal trimer-trimer interactions and a continuous conformational variation between the two trimers related by the scissoring rotation. To characterise the dynamic conformations, we refined two distinct maps representing the extremes of motion: an expanded state—where the trimer-trimer interface is less tight—and a more compact state (Fig. 4B).

**Figure 4.**
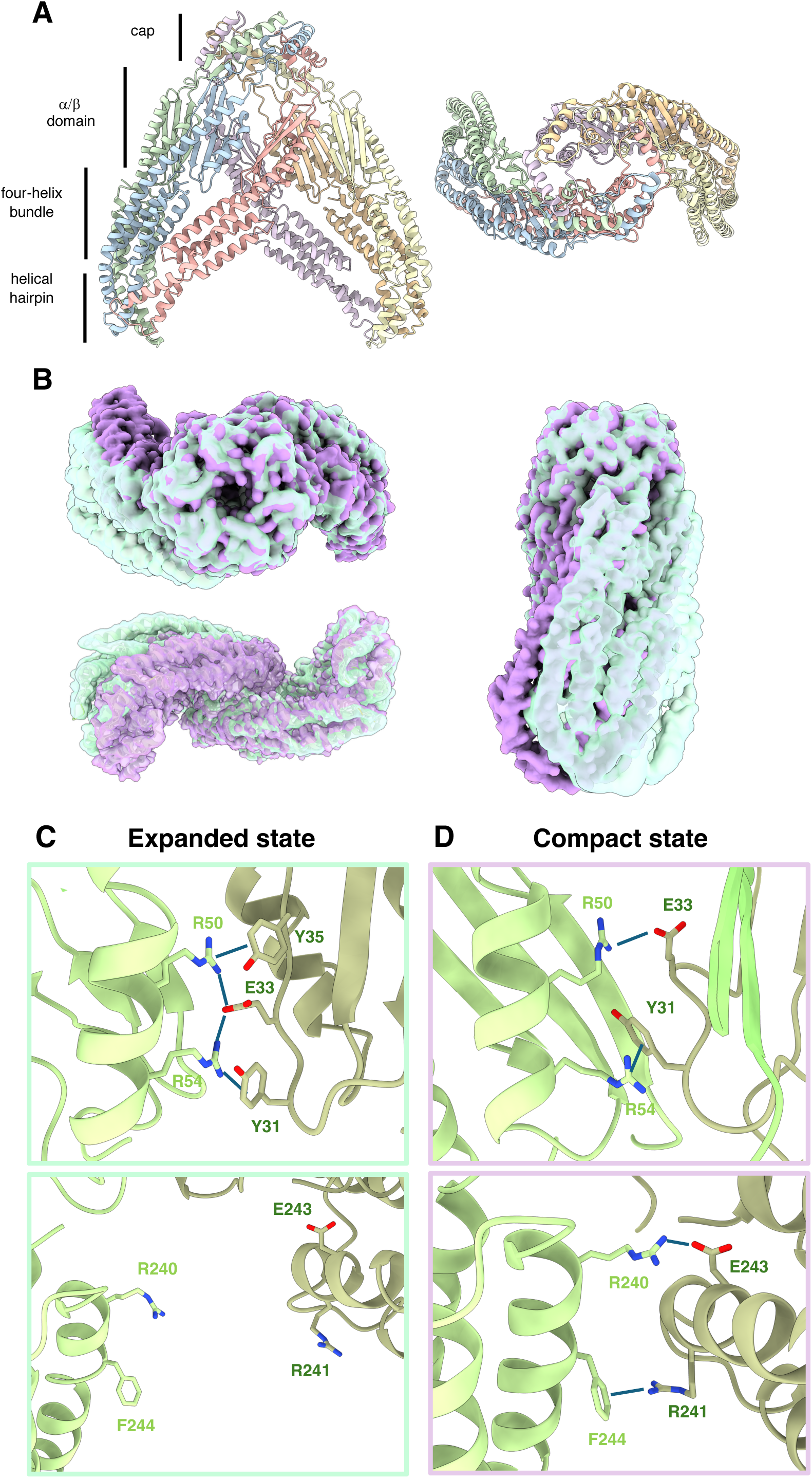
Dynamic trimer-trimer interface stabilises distinct apo-YicC conformations. (A) Annotation of domains and their organisation in the YicC hexamer. (B) Orthogonal views of the superimposed cryoEM maps of the apo-YicC hexamer in the expanded (light green) and compact (purple) states, illustrating the continuous inter-trimer "breathing" motion. (C) Atomic details of the α/β (top) and four-helix bundle domain (bottom) interfaces in the expanded state. Inter-trimer contacts are predominantly maintained by the α/β domain, while the four-helix bundle interface remains open. (D) Corresponding views of the α/β domain, while the four-helix bundle interfaces in the compact state. Hexamer compaction tightens the α/β interface, supported by an electrostatic latch involving E33, R50, and R54. This network is further reinforced by Y31 and Y35, which are positioned to form potential hydrogen bonds and cation-π interactions with the adjacent arginine and glutamate residues. This compaction simultaneously engages a secondary interaction network in the four-helix bundle, where R240, R241, E243, and F244 form additional electrostatic, cation-π, and hydrophobic contacts to lock the trimers together.

In the expanded state, inter-trimer interactions are restricted to the α/β domain (Fig. 4C). These contacts occur between residues in the α-helix of one protomer and the closest adjacent β-strand of the opposing protomer and vice versa. Within this defined-interface, E33, R50, and R54 form an electrostatic “molecular latch”. This network is further reinforced by Y31 and Y35, which are positioned to form hydrogen bonds and cation–π interactions with the adjacent arginine and glutamate residues to stabilise the local architecture. Sequence analysis indicates strong conservation of an acidic residue at position 33 and an arginine at position 50, while position 54 is frequently occupied by positively charged residues, well suited to maintain these inter-protomer contacts even when the hexamer is expanded (Appendix Fig. S2). In the compact state, the interaction network expands beyond the α/β domain to include conserved residues within the four-helix bundle (Fig. 4D). Here, R240 and E243 engage in electrostatic interactions, while F244 from both protomers at the interface is positioned to form hydrophobic packing, π-stacking, and cation–π interactions with R241 that further lock the trimers together. This phenylalanine is highly conserved and a F244A mutation was also shown to impair *E. coli* YicC catalytic activity on short synthetic oligonucleotides (Wu *et al*., 2024). Together, these structures define a dynamic, bipartite inter-trimer latch in *apo*-YicC: conserved α-helix/β-strand engagements tether the expanded state, while additional helical-hairpin domain contacts stabilise the compact state, revealing a breathing mechanism that likely primes the complex for RNA substrate recognition.

### YicC binds an RNA stem loop to position RyhB for cleavage

Having observed the dynamic trimer-trimer interaction in the apo-state of YicC, we next sought to assemble a stable complex with its substrate RyhB for cryoEM analysis. We employed a catalytically deficient YicC mutant, E252A, which was previously shown to significantly impair YicC ribonuclease activity (Huang *et al*., 2023). YicC E252A retains detectable cleavage activity against RyhB but is much less efficient than the wild-type enzyme (Appendix Fig. S5A). Crucially, this variant formed a stable complex with RyhB in our electrophoretic mobility shift assays (EMSAs) (Appendix Fig. S5B), allowing the pre-cleavage state to be trapped. With biochemical validation of the stability of the YicC(E252A)-RyhB complex in solution, we proceeded with cryoEM structure determination. We obtained a map with an overall resolution of 2.7 Å featuring clear density for an A-form RNA helix (Fig. 5A). This cryoEM map allowed us to confidently resolve five YicC protomers and the cap domain of the sixth, while the remainder of the sixth protomer exhibited high flexibility.

**Figure 5.**
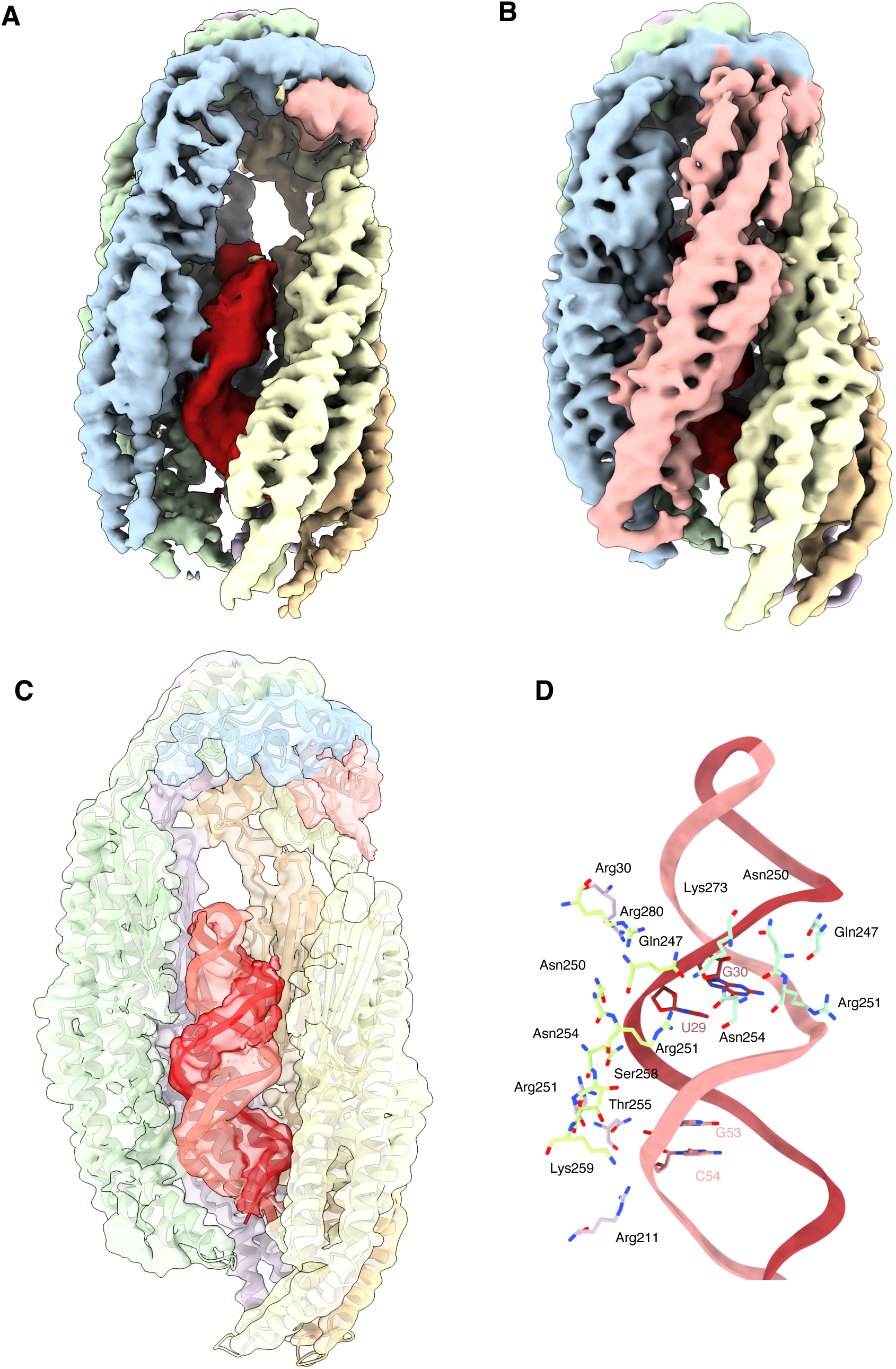
Structure of YicC in complex with RyhB sRNA. (A) CryoEM density of YicC-RyhB_1-95_ (B) CryoEM density corresponding to YicC-RyhB_19-59_ (C) Side view of the final structural model of the YicC-RyhB_19-59_ complex. For clarity, two of the six YicC protomers have been hidden. The electron density corresponding is shown as a transparent surface with the refined atomic model fitted inside. (D) Atomic details of the protein-RNA interface. A network of conserved residues from the retracted trimer makes extensive contacts with the RNA backbone around the U29 and G53 cleavage sites, helping to position the scissile phosphates for catalysis.

The resulting cryoEM model captures how YicC trimers rotate relative to each other upon binding to the RNA substrate, creating an inner cavity that accommodates an RNA hairpin, consistent with recent structural reports (Wu *et al*., 2024). However, our structure reveals a critical difference in substrate orientation. In the previously reported structure of YicC bound to a short synthetic RNA oligo, the loop of the hairpin faces the exterior of the barrel, with the RNA termini buried deep inside the hexameric cavity (Wu *et al*., 2024). By contrast, in our structure with the full-length RyhB substrate, the loop of the RNA hairpin faces towards the cap region of YicC, with the 5’- and 3’-ends of the RNA exiting towards the outside of the barrel. While the overall map resolution was 2.7 Å, the local resolution in the RNA region (> 3 Å) was not sufficient to confidently assign individual RNA bases inside the YicC cavity. However, because the YicC cleavage sites map to the central stem-loop of RyhB (nucleotides 21-58), we hypothesised this to be the captured region. Indeed, the cryoEM density corresponding to the RNA hairpin accommodated an A-form RNA helix modelled from the 38 nucleotides comprising the central RyhB stem-loop.

### YicC adopts an asymmetric conformation to poise the RyhB stem-loop for cleavage

To overcome the conformational flexibility observed with the full-length sRNA, we determined the cryoEM structure of YicC bound to a truncated RyhB fragment encompassing nucleotides 19-59 (RyhB_19-59_), which corresponds to the central RyhB stem-loop. Following dataset classification of the closed, RNA-bound particles (Appendix Fig. S7), we obtained a map at an overall resolution of 3.39 Å where all six YicC protomers enclose the RNA stem-loop (Fig. 5B). The resulting map exhibits clear density for a double-stranded RNA helix, accommodating the central stem-loop of RyhB in the same orientation observed for the full-length YicC-RyhB_1-95_ complex. We modelled the 41-nucleotide RyhB_19-59_ fragment, which fit the RNA density (Fig. 5C).

A notable feature of this RNA-bound YicC hexamer is its marked asymmetry. Upon binding the RyhB substrate, one of the YicC trimers rotates and closes around the RNA, adopting a compacted conformation, whereas the opposing trimer remains relatively extended along the longitudinal axis (Fig. 5B). Next, we examined the local environment surrounding the known cleavage sites (downstream of U29 and G53 in full-length RyhB, corresponding to U11 and G35 in this model). Here we observed that both cleavage sites lie on the same face of the A-form RNA helix and are oriented toward the protomers in the retracted trimer (chains ***a***, ***b***, and ***c***), suggesting that this compacted trimer could act as the primary catalytic hub of the complex (Fig. 5D). Within a 5 Å radius of nucleotides U29 and G30, we observe residues N250, N254, and R273 (from chain ***a***); Q247, N250, R251, N254, and R280 (from chain ***b***); and R30 and Q247 (from chain ***c***). Similarly, within a 5 Å radius of nucleotides G53 and C54, we observe residues T255, S258, and K259 (from chain ***b***), alongside R211, R251, and N254 (from chain ***c***). Because the canonical YicC catalytic residues are not positioned in direct proximity to the scissile phosphates in this model, this structure likely captures a stalled, pre-cleavage state. In this state, the dense network of polar and basic interactions could anchor the substrate and precisely orient the RNA backbone, poising the scissile phosphates for the final conformational shift required for nucleophilic attack. With the exception of N250, these orienting residues are strictly conserved across bacterial YicC homologues (Appendix Fig. S2). The positioning of these conserved residues along one face of the RNA helix provides a structural basis for how YicC would specifically engage and poise highly structured RNA targets for cleavage.

In all apo and RNA-bound structures, the α/β domain, four-helix bundle, and helical hairpin form an effectively rigid unit that is invariant with the scissoring rotation, and moves as a body with respect to the hexameric cap. Notably, the helical hairpin packs in a markedly non-equivalent manner within each unit, so that the third protomer adopts a bent helical arrangement that packs to form a local four-helix bundle with the central protomer (Fig. 6). In the structure of the YicC-RyhB_1-95_ complex, where there are only 5 YicC protomers that are well packed, the helical hairpin is in equilibrium between two packing modes for one of the protomers (Fig. 6C). If the conformation of the invariant trimeric unit is represented as protomers being in states **abc**—where **c** denotes a protomer where the helical hairpin is in a bent conformation—, then the 5 protomers in the YicRyhB_1-95_ complex are in equilibrium between configurations **ababc** and **abcbc**. This conformational switch from **a** to **c** brings the helical hairpin of the **c/a** subunit into proximity of the embedded RNA stem-loop (Fig. 6C).

**Figure 6.**
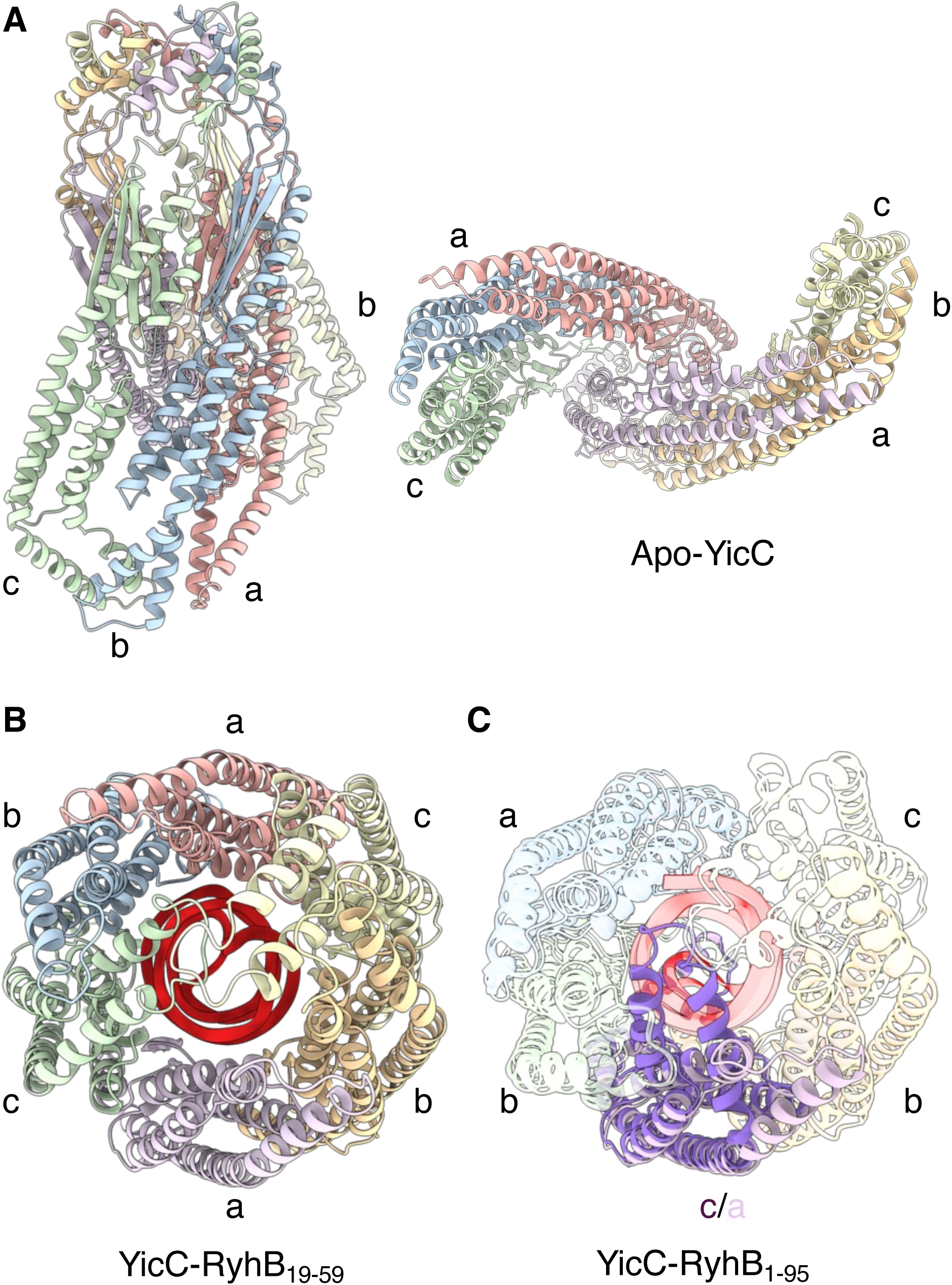
The invariant abc trimer and permutation of tertiary conformational states in the YicC-RyhB_1–95_ complex. The **abc** trimeric unit in the apo state (A) and in the RNA-bound state with RyhB_15-59_ (B). The three conformational states for the helical hairpin in YicC protomers are labelled as **a**, **b** and **c**. (C) Conformational switch of the helical hairpin domain of YicC in the complex with RyhB_1-95_. Models corresponding to YicC-RyhB_1-95_ in configurations **abcbc** and **ababc** are superimposed. Protomer that shows the conformational switch is highlighted with the RNA molecule and the rest of protomers dimmed. Only five out of six protomers are visible as only the cap domain could be modelled for the sixth protomer due to its flexibility.

Collectively, these findings support a model where the *apo*-YicC hexamer exists in a stable, latched conformation that could act as an "RNA scanning" or "surveillance" complex. In this state, the enzyme is structurally poised for substrate recognition but remains catalytically inactive. The binding of a specific RNA structural motif would trigger a signal that breaks the electrostatic and hydrophobic latches at the trimer-trimer interface, inducing an asymmetric conformational change into a cleavage-competent state. This regulatory mechanism, dependent on a highly conserved set of interface residues, appears to be a fundamental feature for maintaining enzymatic fidelity and preventing spurious RNA degradation.

## Discussion

YicC is a widely conserved but mechanistically enigmatic ribonuclease whose precise biological role in bacterial RNA processing remains to be fully defined. Despite recent structural insights into the hexameric assembly of the apo- and RNA-bound enzyme, the mechanisms governing its conformational dynamics, substrate recognition, and catalytic activation have remained unclear. In this study, we determined the cryoEM structures of *E. coli* YicC in both its dynamic *apo* state and its physiologically relevant, RNA-bound state. Our structural analysis reveals that *apo*-YicC is not a static open barrel but rather exists in a continuous "breathing" equilibrium mediated by a conserved bipartite interaction network at the trimer-trimer interface. Furthermore, by capturing YicC in complex with the physiological RyhB sRNA, we resolved the architecture of its central stem-loop bound within the central cavity. We demonstrate that the binding of this RNA element induces a structural asymmetry in the homo-hexameric ribonuclease. This asymmetric compaction orients the RNA stem-loop into a biologically viable conformation—distinct from previously reported structures with short synthetic oligos—but also organises a dense network of conserved residues that poises the scissile phosphates for cleavage. Together, these findings provide a comprehensive structural basis for how YicC acts as a dynamic surveillance complex that requires specific, structured RNA triggers for catalytic activation.

A previous study positioned YicC as a putative adaptor for the 3’-5’ exoribonuclease PNPase, based on the observation that YicC-mediated reduction of RyhB levels in *E. coli* was lost in a *pnp* deletion strain (Chen *et al*., 2021). Furthermore, different studies using bacterial two-hybrid assays have shown that PNPase can interact *in vivo* with *E. coli* YicC, and its homologues in *B. subtilis* and *C. difficile* (Chen *et al*., 2021; Ingle *et al*., 2022; Martins *et al*., 2025). Our biochemical results, however, indicate that purified YicC can readily cleave its biological substrate, RyhB, *in vitro* in the absence of PNPase. We propose that the previous *in vivo* observations can be explained by the dual role of PNPase, which also acts as a protective chaperone for certain sRNAs (Bandyra *et al*., 2016; Dendooven *et al*., 2021). In a *pnp* deletion strain, inherently low basal levels of RyhB likely create a threshold effect that could mask any further YicC-mediated degradation. Using a fluorescent reporter system, we confirmed that YicC directly cleaves RyhB in a PNPase-independent manner.

Our cleavage assays show that YicC exhibits distinct modes of activity depending on the presence of the RNA chaperone Hfq (Fig. 1). While YicC degrades unprotected RyhB completely, the presence of Hfq restricts YicC to processing RyhB in a highly specific manner, generating two discrete cleavage products by cutting at U29 and G53 within the central stem-loop. This suggests that Hfq remodels RyhB’s conformation, exposing defined positions within a structured region for YicC to recognise and cleave. Furthermore, YicC exhibits substrate selectivity. This ribonuclease efficiently cleaves the Class I sRNA RyhB and the intermediate Class I/II sRNA RprA, but fails to degrade other tested sRNAs, including MicA, ArcZ (ClassI), and CyaR (Class II). This selectivity could extend beyond the identity of the sRNA to its functional state. We found that RNA-RNA base-pairing—whether through a canonical mRNA target (such as *csgD*) or an RNA sponge (such as ETS^LeuZ^)—protects the sRNA from YicC-mediated degradation. Because the YicC hexameric cavity is structurally tailored to accommodate a specific RNA hairpin architecture, the formation of an extended RNA duplex likely creates severe steric clashes or occludes necessary recognition motifs. This suggests a refined physiological role for YicC: acting as a clearance factor that selectively degrades excess or orphaned sRNAs without prematurely dismantling active regulatory complexes.

### A dynamic trimer-trimer interface stabilises the open apo conformation

To understand the structural basis for this substrate specificity, we determined the solution-state structure of the apoenzyme. While previous crystal structures captured static open conformations (Wu et al., 2024; Huang et al., 2023), our cryoEM analysis reveals that *apo*-YicC in solution is highly dynamic, existing in a continuous "breathing" equilibrium between an expanded and a compact state. This motion is governed by a conserved, bipartite inter-trimer latch. In the expanded state, the hexamer is tethered primarily by an α/β domain electrostatic latch, driven by residues such as E33, R50, and R54, and reinforced by adjacent tyrosine residues. As the hexamer "breathes" into the compact state, a secondary interaction network engages, further locking the trimers together through additional electrostatic and cation-π interactions centred around the strictly conserved F244, within the four-helix bundle domain.

We hypothesise that this dynamic, bipartite latching mechanism stabilises the hexamer in a secure "surveillance" conformation that would allow the enzyme to continuously sample cellular RNAs. Because the YicC cavity is accessible in this breathing state, the enzyme can likely bind non-substrate RNAs; however, these off-target molecules would fail to provide the necessary conformational cues to overcome the energy barrier of the inter-trimer latch. Only the engagement of a *bona fide* substrate with the correct structural architecture—such as the central stem-loop of RyhB—could break these distinct domain latches. This specific recognition event would drive the hexamer out of its auto-inhibited equilibrium and into the catalytically active conformation, thereby ensuring fidelity and preventing the spurious degradation of off-target RNAs.

### Substrate identity dictates binding orientation and catalytic asymmetry

Upon binding the central stem-loop of RyhB, the inter-trimer latches are released, and YicC transitions to a closed conformation. Consistent with the RNA-bound state reported by Wu *et al*. (2024), YicC adopts an asymmetric conformation upon substrate binding, with one trimer rotating to close upon the RNA hairpin (the "retracted" trimer) while the other remains relatively extended (Wu *et al*., 2024). Notably, our structure reveals a distinct mode of substrate binding compared to this previous model. In the structure reported by Wu *et al*. (2024) using a short, synthetic oligonucleotide, the hairpin loop is oriented towards the exterior of the complex, with the 5′ and 3′ termini buried inside the central cavity (Wu *et al*., 2024). In contrast, our structure with RyhB substrate shows the opposite orientation: the hairpin loop is positioned deep within the cavity facing the hexamerisation cap, while the 5′ and 3′ ends of the RNA are directed towards the exterior of the barrel (Fig. 5). This orientation would allow a stem-loop from a full-length sRNA to enter and exit the enzymatic cavity without severe steric clashes. Mapping the cleavage sites of RyhB onto this architecture, suggests that cleavage originates from the retracted trimer. Furthermore, the positioning of the RyhB substrate in our map places the previously proposed catalytic triad (E216, E217, E281) at a significant distance from the scissile phosphates. This suggests our structure captures a stable, stalled pre-cleavage state where the RNA is securely bound and functionally oriented, prior to the final conformational engagement of the catalytic machinery.

### Evolutionary implications

The observed structural specificity presents an important evolutionary consideration. While the YicC protein is broadly conserved across the bacterial domain, its substrate, RyhB, is phylogenetically restricted to enterobacteria and specific *Vibrio* species (Oglesby-Sherrouse & Murphy, 2013). The discrepancy between the broad conservation of the enzyme and the narrow distribution of its known substrate strongly implies that YicC function is not exclusively linked to RyhB. A recent report identified a small RNA, SQ528, as a putative target of the YicC homologue in *C. difficile* (Martins *et al*., 2025). We propose that YicC orthologues have co-evolved with RNA substrates based on conserved 3D structural features rather than primary sequence identity. The identification of native YicC substrates in a broader range of species through future transcriptomic approaches will be necessary to fully understand how this ancient ribonuclease has been adapted to distinct regulatory networks across bacterial lineages.

## Materials and methods

### Bacterial strains and plasmids

*E. coli* strains, plasmids used in this study are listed in Appendix Tables S1 and S2, respectively. Primers used in this study were ordered from Sigma-Aldrich and are listed in Appendix Table S3. The *sodB* reporter was adapted from a fluorescent construct described by Chen *et al*. (2021). A *sodB-mCherry* translational fusion under a constitutive promoter (cp26) regulation was PCR amplified from a synthetic gene and cloned into a pZS21 vector. pBAD-RyhB was constructed by cloning the RyhB sequence under the cp26 promoter into a pBAD vector. pZA31-YicC vectors were engineered by PCR amplifying YicC from *E. coli* MG1655 gDNA and cloned into pZA31 under the cp26 promoter.

### Protein preparations

#### Hfq

Hfq was expressed from pET-Duet 1 vector in BL21(DE3) cells at 37 °C, induced with 1 mM IPTG at OD_600_ = 0.4-0.6 and harvested by centrifugation 3-4 hours after induction. Cells were resuspended in Hfq Lysis Buffer (50 mM Tris-HCl pH 8.0, 1.5 M NaCl, 250 mM MgCl_2_, cOmplete™ EDTA-free Protease Inhibitor Cocktail [Roche]. Resuspended cells were lysed using an Emulsiflex C5 (Avestin) high-pressure homogeniser (1000 bar). The lysate was clarified by centrifugation (4 °C, 37500 g, 30 mins), and the supernatant was incubated at 85 °C for 20 minutes and centrifuged (20 °C, 37500 g, 30 mins). The resulting supernatant was subjected to ammonium sulfate precipitation (1.5 M) and centrifuged (20 °C, 37500 g, 30 mins). The supernatant was filtered (Sartorius Minisart® 0.45 µm) and loaded on 5 mL HiTrap Butyl HP (Cytiva), previously equilibrated with Butyl Buffer A (50 mM Tris pH 8.0, 1.5 M NaCl, 1 M (NH_4_)_2_SO_4_). A gradient of Butyl Buffer B (50 mM Tris pH 8.0) was used for elution. Fractions containing Hfq were pooled and diluted 1:5 with Butyl Buffer B. Pooled sample was then loaded on 5 mL HiTrap Heparin HP (Cytiva), previously equilibrated with Heparin Buffer A (50 mM Tris-HCl pH 8.0, 100 mM NaCl, 100 mM KCl), and eluted with a gradient of Heparin Buffer B (50 mM Tris-HCl pH 8.0, 1 M NaCl, 100 mM KCl). Fractions containing Hfq were pooled and concentrated (Amicon® Ultra-15 30 KDa, Millipore). Protein solution was loaded on Superdex 200 Increase 10/300 GL size exclusion column (S200; Cytiva) previously equilibrated with Hfq Storage Buffer (50 mM Tris-HCl pH 8.0, 100 mM NaCl, 100 mM KCl, 5 % glycerol). Fractions from S200 were evaluated by SDS-PAGE and concentration of Hfq was determined using a NanoDrop™ One spectrophotometer (Thermo Fisher Scientific).

#### YicC

The DNA corresponding to *E. coli* YicC was PCR amplified from MG1655 genomic DNA using primers YicC_TEV_Fw and YicC_st_Rv (Appendix Table S3) and cloned into pExp-His-xMBP-TEV vector. The E252A mutation was introduced by PCR-based mutagenesis using YicC_E252A_Fw and YicC_E252A_Rv primers. The fusion protein, carrying N-terminal 8xHis tag, followed by maltose-binding protein (MBP) and TEV protease cleavage site preceding YicC was expressed in Evo21(DE3) cells during overnight induction with 0.4 mM IPTG at 18°C. After cell disruption using Emulsiflex C5 (Avestin) homogeniser, the protein was first purified by IMAC using Ni-NTA resin (Qiagen) and then by affinity chromatography using amylose resin (NEB). The His-MBP-fusion tag was cleaved off by TEV protease (produced in-house) for 16 h at 18°C and removed by passing the protein solution through Ni-NTA column. The protein was further purified by anion-exchange chromatography using HiTrap Q HP column (Cytiva) and then by heparin affinity chromatography using HiTrap Heparin HP column (Cytiva). Finally, the protein was purified by size-exclusion chromatography using Superose 6 Increase 10/300 column (Cytiva) equilibrated in GF buffer (20 mM HEPES-NaOH pH 7.2, 200 mM NaCl, 2 mM MgCl_2_, 0.5 mM TCEP). The protein concentration was adjusted to 3.1 mg/ml (15 µM for hexamer) using Amicon Ultra concentrator with 30 kDa cut-off (Millipore).

#### PNPase

*E. coli* PNPase was prepared as described in (Dendooven *et al*, 2021; Paris *et al*, 2025). In brief, *E. coli* BL21(DE3) cells were transformed with PNPase expression plasmid, grown in 2xYT media at 37°C and induced at early exponential phase with 0.5 mM IPTG, and then the temperature was reduced to 25°C. After four hours, cells were harvested by centrifugation, suspended in Lysis Buffer 1 (20 mM Tris-HCl pH 8.0, 150 mM NaCl, 150 mM KCl, 5 mM MgCl2, 1 mM EDTA), lysed using a cell disruptor (Emulsiflex, Avestin, 1000 bar), and the lysate was clarified by centrifugation (4°C, 37500g, 30 mins). Ammonium sulfate was added to the supernatant to 51.3% saturation. Following centrifugation (4°C, 37500g, 30 mins), the pellet was resuspended in Q Buffer A’ (20 mM Tris-HCl pH 8.5, 0.5 mM TCEP, 10% v/v glycerol, EDTA-free protease inhibitor cocktail (Roche)) to solubilise PNPase. The solubilised sample was loaded onto a 5 mL HiTrap Q column (GE Healthcare), equilibrated in Q Buffer A (20 mM Tris-HCl pH 8.5, 30 mM NaCl, 0.5 mM TCEP, 10% v/v glycerol) and eluted with a gradient of Q Buffer B (20 mM Tris-HCl pH 8.5, 1M NaCl, 0.5 mM TCEP, 10% v/v glycerol). Fractions containing PNPase were pooled, mixed with an equal volume of Supplementation Buffer (1 mM MgCl_2_, 45 mM NaH_2_PO_4_ pH 7.9, 0.9 M (NH_4_)_2_SO_4_, 1 mM TCEP), and loaded on a 5 mL HiTrap butyl-Sepharose HIC column (GE Healthcare) equilibrated with BS Buffer A (50 mM Tris-HCl pH 7.5, 1 M (NH_4_)_2_SO_4_, 0.5 mM TCEP) and eluted with a gradient of BS Buffer B (50 mM Tris-HCl pH 7.5). Fractions enriched in PNPase were pooled, concentrated, and loaded on a Superdex 200 Increase 10/300 gel filtration column equilibrated with PNPase Storage Buffer (20 mM Tris-HCl pH 8.0, 150 mM NaCl, 5 mM MgCl_2_, 0.5 mM TCEP, 10% (v/v) glycerol).

#### RNase E (1-598)

The *E. coli* RNase E (1-598) construct was PCR amplified using the *rne* gene as a template and primers RNase_E_1-598_Fw and RNase_E_1-598_Rv (Appendix Table S3), and cloned into a pET15 vector. BL21(DE3) cells, transformed with RNase E (1-598) expression plasmid, were grown in 2×YT media at 37°C and induced with 1 mM IPTG at OD_600_=0.6. RNase E (1-598) was expressed for 3h at 37°C. Following protein expression, cells were harvested and resuspended in NiA buffer (50 mM Tris-HCl pH 7.5, 1 M NaCl, 100 mM KCl, 5 mM MgCl_2_, 20 mM Imidazole pH 7.5, 0.5 mM TCEP, 0.02% w/v β-DDM, complete EDTA-free protease inhibitor tablet (Roche)), supplemented with 1% TritonX and 0.02% w/v β-DDM, and subsequently disrupted using Emulsiflex C5 (Avestin) homogeniser. The lysate was supplemented with DNase I (2 µg/ml), clarified by centrifugation (4°C, 30 minutes, 25000 g) and purified by Immobilized Metal Ion Affinity Chromatography (IMAC) using Ni-NTA resin (Qiagen) equilibrated with NiA buffer. The His-tagged RNase E (1-598) was eluted in buffer NiB (50 mM Tris-HCl pH 7.5, 1 M NaCl, 100 mM KCl, 5 mM MgCl_2_, 500 mM Imidazole pH 7.5, 0.5 mM TCEP, 0.02% w/v β-DDM) and diluted 3-fold with SP A buffer (50 mM Tris-HCl pH 7.5, 50 mM NaCl, 10 mM KCl, 0.5 mM TCEP 0.02% w/v β-DDM). The diluted sample was loaded on a 1 ml HiTrap SP HP column (GE Healthcare), equilibrated with SP A buffer. RNase E (1-598) was eluted with a linear gradient 0-100% of SP B buffer (50mM Tris-HCl pH 7.5, 2 M NaCl, 10 mM KCl, 0.5 mM TCEP, 0.02% w/v β-DDM), concentrated and loaded onto a Superdex200 Increase gel filtration column equilibrated with S200 buffer (50 mM HEPES pH 7.5, 400 mM NaCl, 100 mM KCl, 5 mM MgCl_2_, 0.5 mM TCEP, 0.02% w/v β-DDM).

### *In vitro* transcription (IVT)

The sRNAs RprA, MicA, CyaR, RyhB and all RyhB variants were transcribed *in vitro* using recombinantly expressed and purified T7 RNA polymerase. For MicA, CyaR, RyhB and all RyhB variants, reverse complimentary oligos were heat-annealed to form a dsDNA template that included a promoter region for the T7 polymerase (sequences available in Appendix Table S3). Briefly, 20 μM of each oligo were mixed in heat-annealing buffer (50 mM Tris-HCl pH 7.5, 10 mM MgCl_2_, 5 mM DTT, 0.1 mM spermidine), incubated for 15 min at 95°C and allowed to cool to room temperature. DNA templates for ArcZ, RprA and *csgD* (nucleotides -147 to

+18 relative to the translation start site) were generated by PCR from *E. coli* K12 MG1655 gDNA to include an upstream T7 promoter (sequences of primers available in Appendix Table S3). PCR products were cleaned up using spin tubes carrying silica-membranes (QIAquick PCR purification kit), loaded on a 0.8 % agarose gel and gel purified. 100 μl IVT reactions containing 1 mM of template dsDNA, T7 RNA polymerase (0.25 mg/mL), 5 mM of each rNTP, 10 mM DTT, 0.5 U/μL RNaseOUT™ Recombinant Ribonuclease Inhibitor [Invitrogen] and 3% v/v DMSO were incubated in IVT reaction Buffer (40 mM Tris-HCl pH 8, 25 mM MgCl_2_, 2 mM spermidine) at 37°C for 3 h. IVT products were incubated with Turbo DNase I (Invitrogen) at 37°C for 30 minutes. DNase treated samples were incubated at room temperature with 50 mM EDTA for 5 mins. RNA was cleaned up using PureLink™ RNA Micro Kit (Invitrogen) columns, checked on 8 % urea-PAGE and stored in nuclease-free water at -20°C.

### RNA Degradation assays

YicC cleavage activity was tested using different RNA as substrates in the presence or absence of PNPase and/or Hfq. RNA was annealed by incubating for 2 min at 50°C and letting cool to room temperature. 0.2 μM RNA was incubated at 37°C in RNA cleavage buffer (20 mM Tris pH 7.5, 100 mM NaCl, 1 mM DTT, 10 mM MgCl_2_, 2 mM sodium phosphate) in the presence or absence of equimolar concentrations of Hfq. A time point 0 was taken immediately before enzyme addition and reactions were started by adding YicC and or PNPase. The reactions were stopped at the indicated time points (1, 5, 15 min) by adding an equal volume of 0.5 mg/ml Proteinase K in Proteinase K buffer (100 mM Tris-HCl pH 7.5, 150 mM NaCl, 12.5 mM EDTA and 1% w/v SDS) and incubated at 50°C for 30 min. Products from each timepoint were resolved on a 10 % polyacrylamide, 7.5M urea gel in 1xTBE, stained in SYBR Gold solution (Invitrogen) and visualised by fluorescence imaging.

### Rapid amplification of cDNA ends (RACE)

YicC cleavage sites on RyhB were identified using 5’ RACE. A 20 μL reaction sample was prepared by annealing RyhB (200 nM) as explained above and incubating it in the presence of equimolar concentration of Hfq at 37°C in RNA cleavage buffer. YicC was added at 200 nM and the reaction was quenched after 15 min by adding 20 mM EDTA. The sample was cleaned up using Monarch^®^ Spin RNA Cleanup Kit (NEB). A DNA oligo including a NcoI restriction site (sequences available in Appendix Table S3) was ligated to the 5’ end of the cleavage products using T4 RNA ligase 1 (NEB) following the manufacturer’s protocol: cleaned up cleavage products were incubated at 25°C for 2 h in T4 RNA Ligase Reaction Buffer (50 mM Tris-HCl, pH 7.5, 10 mM MgCl_2_, 1 mM DTT) supplemented with 15 % (wt/vol) PEG 8000, 10 % DMSO, 0.5 U/μL RNaseOUT^™^ Recombinant Ribonuclease Inhibitor [Invitrogen], 1 mM ATP, 1 μl (10 units) T4 RNA Ligase 1 and 50 pmol of DNA oligo in a 20 μL reaction volume. Ligation product was cleaned up using an RNA clean up kit and used as template for reverse transcription using the SuperScript IV First-Strand Synthesis System (Invitrogen) according to the manufacturer’s protocol. The reaction mixture was incubated at 55°C for 10 min, followed by inactivation at 80°C for 10 minutes and RNase H treatment at 37°C for 20 min. 2 μL of cDNA served as template for PCR reaction using Phire Hot Start II DNA polymerase (Thermo Scientific). In addition to the primer including a NcoI restriction site mentioned above, a second primer with a HindIII restriction site followed by a sequence reverse-complementary to the 3’ end of RyhB was used for the amplification (sequences available in Appendix Table S3). Following the PCR, samples were cleaned up using GeneJET PCR Purification Kit (Thermo Scientific), digested with NcoI and HindIII (NEB), and cloned into a NcoI/HindIII pre-digested pET-Duet 1 vector using T4 DNA ligase (NEB). Ligation products were transformed into chemically competent *E. coli* DH5α. 16 colonies were grown in liquid culture overnight and vectors extracted by miniprep, using GeneJET Plasmid Miniprep Kit (Thermo Scientific), were sent for Sanger sequencing. The same 5’ RACE protocol was followed with a sample were YicC was not added as a control of RyhB integrity.

### RNA binding assays

Stability of YicC-RyhB complexes was tested by electrophoretic mobility shift assays (EMSA). RyhB was annealed by incubating for 2 min at 50°C and letting cool down to room temperature. 10 μL samples were prepared by incubating 50 nM annealed RyhB in binding buffer (20 mM Tris pH 8.0, 150 mM KCl, 2.5 mM MgCl_2_, 1 mM TCEP) at 30 °C for 20 min with YicC E252A (catalytically inactive), in the presence or absence of Hfq. Samples were transferred to ice, supplemented with 5 μL of 50 % glycerol in binding buffer and loaded on a 6% (wt/vol) native polyacrylamide gel. The gel was run on TBE buffer for 80 min at 120 V and imaged on a Syngene InGenius3 system. The concentrations of YicC and Hfq used are indicated on figures.

### CryoEM sample preparation

The interaction between YicC and RyhB was structurally characterised using cryo-EM. RyhB was annealed by incubating for 2 min at 50°C and letting cool down to room temperature. Subsequently, annealed RyhB (nucleotides 1-95 or 19-59) was incubated with YicC E252A at equimolar concentrations (12.5 µM) in binding buffer to form complexes. Samples were also prepared for YicC E252A at 15 µM in the absence of RyhB (Apo-YicC). 3 µL of each sample was applied onto UltrAufoil^®^ R1.2/1.3 grids (Quantifoil) previously glow-discharged (PELCO easiGLOW: 60 s, 25 mA, 0.39 mBar). Grids were blotted, and sample vitrified in liquid ethane using a FEI Vitrobot (IV) (100% humidity, 4°C, blotting force 0, blot time 3 sec).

### CryoEM data collection

18,570 multi-frame movies for an apo-YicC dataset were collected at the UK national electron Bio-Imaging Centre (eBIC) on a Titan Krios transmission electron microscope operated at 300 kV and equipped with a Gatan K3 detector and energy filter using EPU software (Thermo Fisher Scientific). For datasets corresponding to YicC-RyhB complexes, 9,869 (first YicC-RyhB_1-95_ dataset), 10,636 (second YicC-RyhB_1-95_ dataset) and 10,329 (YicC-RyhB_19-59_) multi-frame movies were collected at the School of Biological Sciences Cryo-EM Facility University of Cambridge, on a Titan Krios G3 (Thermo Fisher Scientific) operated at 300 kV and equipped with a Falcon 4i detector and SelectrisX energy filter (Thermo Fisher Scientific) set to a 10 eV slit width, using EPU software (Thermo Fisher Scientific). The data collection parameters are summarised in Appendix Table S4.

### CryoEM data processing

#### Apo-YicC

For apo-YicC, movies were motion-corrected and dose-weighted using RELION’s implementation of the MotionCor2 algorithm, and CTF parameters were estimated on the resulting micrographs using the RELION wrapper to CTFFIND 4.1 (Rohou and Grigorieff, 2015; Scheres, 2012; Zheng et al., 2017). A schematic summary of the processing workflow can be found in Appendix Fig. S6. A particle set was picked using Topaz pre-trained model for automated picking (Bepler *et al*, 2019). Following extraction, particles were subjected to a round of reference-free 2D classification using the VDAM algorithm (Kimanius *et al*., 2021). A subset of 752,660 particles from high-quality 2D classes was imported into cryoSPARC v4.7.0. and used to generate an *ab initio* model (Punjani *et al*., 2017). These particle set was further curated via heterogeneous refinement. To explore the conformational landscape, we performed 3D Variability Analysis (3DVA) using the "cluster" mode. Because the motion appeared continuous, we selected particle subsets corresponding to the two extreme conformational states for subsequent non-uniform refinement and local refinement. The resulting classes—150,108 particles in the expanded state and 215,534 in the compact state—were subjected to non-uniform local refinement to improve alignment and map resolution.

#### YicC-RyhB_1-95_

For YicC-RyhB_1-95_, motion correction, CTF estimation particle picking and extraction were carried out in Warp 0.9 (Tegunov & Cramer, 2019). Subsequently particles were imported into CryoSPARC v 4.7.0 (Punjani *et al*, 2017). A schematic summary of the processing workflow can be found in Appendix Fig. S7. Briefly, a round of 2D classification was performed to remove erroneous picks. Classes presenting YicC in different orientations were used to generate *ab initio* models. These sets were curated via heterogeneous refinement to select particles corresponding closed RNA-bound YicC. The particle set was then subjected to two rounds of 3D classification using a focus mask corresponding to the inner cavity where the RNA is embedded. Particle sets with a higher occupancy in the RNA cavity were selected for a final Non-Uniform refinement to obtain the final map.

#### YicC-RyhB_19-59_

For YicC-RyhB_19-59_, motion correction, CTF estimation particle picking and extraction were carried out in Warp 0.9 (Tegunov & Cramer, 2019). Subsequently particles were imported into CryoSPARC v 4.7.0 (Punjani *et al*, 2017). A schematic summary of the processing workflow can be found in Appendix Fig. S8. Briefly, 2D classification was performed to remove erroneous picks. Classes showing YicC in a closed conformation in different orientations were used to generate *ab initio* models and subsequently curated via heterogeneous refinement. Following heterogeneous refinement, one class showed YicC in a closed conformation and with clear density for the RNA and was refined via non-uniform refinement. Focused 3D classification was performed using a mask excluding the flexible protomer of YicC, resulting in two classes with improved density for the RNA. These two classes were used for subsequent rounds of particle curation and 3D refinement to obtain the final map.

### Model building and refinement

The resolutions of the maps were estimated by 0.143 cut-off of the Fourier Shell Correlation (FSC) from two independently refined half-reconstructions. The final consensus reconstruction for each dataset was used as input for automated model building using ModelAngelo (Jamali *et al*., 2024). For YicC-RyhB_19-59, the 41-nucleotide sequence for the stem-loop was modelled using AlphaFold 3 and docked into the cryoEM density using rigid body fit in ChimeraX 1.18 (Pettersen *et al*., 2004). Gaps in the model obtained from ModelAngelo were manually built using Coot 0.9 (Emsley *et al*, 2010). The completed model was then refined against the cryo-EM density using phenix.real_space_refine (Afonine *et al*., 2018). Secondary structure restraints were applied to the RNA bases forming the stem and to residues forming alpha helices and beta-strands in YicC. Ramachandran outliers were corrected using ISOLDE 1.6 (Croll, 2018). Finally, the quality of the stereochemistry was evaluated using the PDB validation tool in Phenix 1.21 (Afonine *et al*., 2018).

### Phylogenetic analysis

A base phylogenetic tree representing major bacterial lineages at the phylum level was downloaded in Newick format from AnnoTree, which uses the Genome Taxonomy Database (GTDB) Release 214 topology (Mendler *et al*., 2019; Parks *et al*., 2018). To enhance visual clarity and ensure robust representation, this tree was filtered using the ape package in RStudio to retain only phyla containing a minimum of 10 sequenced genomes (Paradis & Schliep, 2019). This filtering step mathematically preserved the underlying evolutionary topology and branch lengths (expected substitutions per site) of the remaining lineages. The proportion of genomes encoding a *YicC* homologue within each phylum was calculated and mapped onto the filtered phylogeny. The final figure was visualised and annotated using the Interactive Tree of Life (iTOL) web platform (Letunic & Bork, 2024). Within the visualisation, external colour gradients designate the percentage of *YicC* prevalence, while internal branch colours were manually curated to infer the ancestral presence (red) or absence (black) of the gene across clades.

A protein sequence logo was built to represent the sequence conservation of YicC-like proteins in Bacteria. 87 homologues of *E. coli* YicC were identified across Bacteria by BLASTP searches against the NCBI nr database, using the BLOSUM62 substitution matrix, or BLOSUM45 to enhance detection of distant homologues within highly divergent bacterial taxa. Representative sequences were aligned with MAFFT v7 employing the L-INS-i algorithm and BLOSUM62 scoring matrix to maximise alignment accuracy (Katoh & Standley, 2013). The alignment was subjected to automated trimming with trimAl v1.4.1 (gap threshold = 0.2) to remove poorly aligned or gap-rich regions that may confound phylogenetic inference (Capella-Gutiérrez *et al*., 2009). The curated multiple sequence alignment was used to generate a sequence logo with WebLogo 3 (Crooks *et al*., 2004).

## Acknowledgements

The work was supported by Wellcome Trust Investigator Award to BFL (222451/Z/21/Z). GP is supported by a Levy studentship. KKG is supported by a Boehringer Ingelheim Fonds PhD studentship. We thank Steven W. Hardwick, Katarzyna J. Bandyra, Kai Pappenfort, Eyal Maori and Yanjie Chao for stimulating and helpful discussion discussions. We thank the School of Biological Sciences Cryo-EM facility at the Department of Biochemistry, University of Cambridge for their support with cryoEM data acquisition. We acknowledge Diamond for access and support of the cryo-EM facilities at the UK national electron Bio-Imaging Centre (eBIC), proposal BI44655.

## Data availability

The refined coordinates and maps have been deposited with the PDB. The access codes for apo-compact state model and map are 29ZL and EMD-57472; for the apo-expanded state, the codes are 30AB and EMD-57480; for YicC/RyhB_1-95_ with first data set with 100000 particles in state 1, the codes are 30BE and EMD-57534; for YicC/RyhB_1-95_ with second data set of 201000 particles, in state 1, the codes are 30EK and EMD-57665; for YicC/RyhB_1-95_ with second data set of 201000 particles, in conformational state 2, the codes are 30ET and EMD-57665; for YicC/RyhB_19-59_, the codes are 30FI and EMD-57694.

## Appendix

**Appendix Figure S1.**
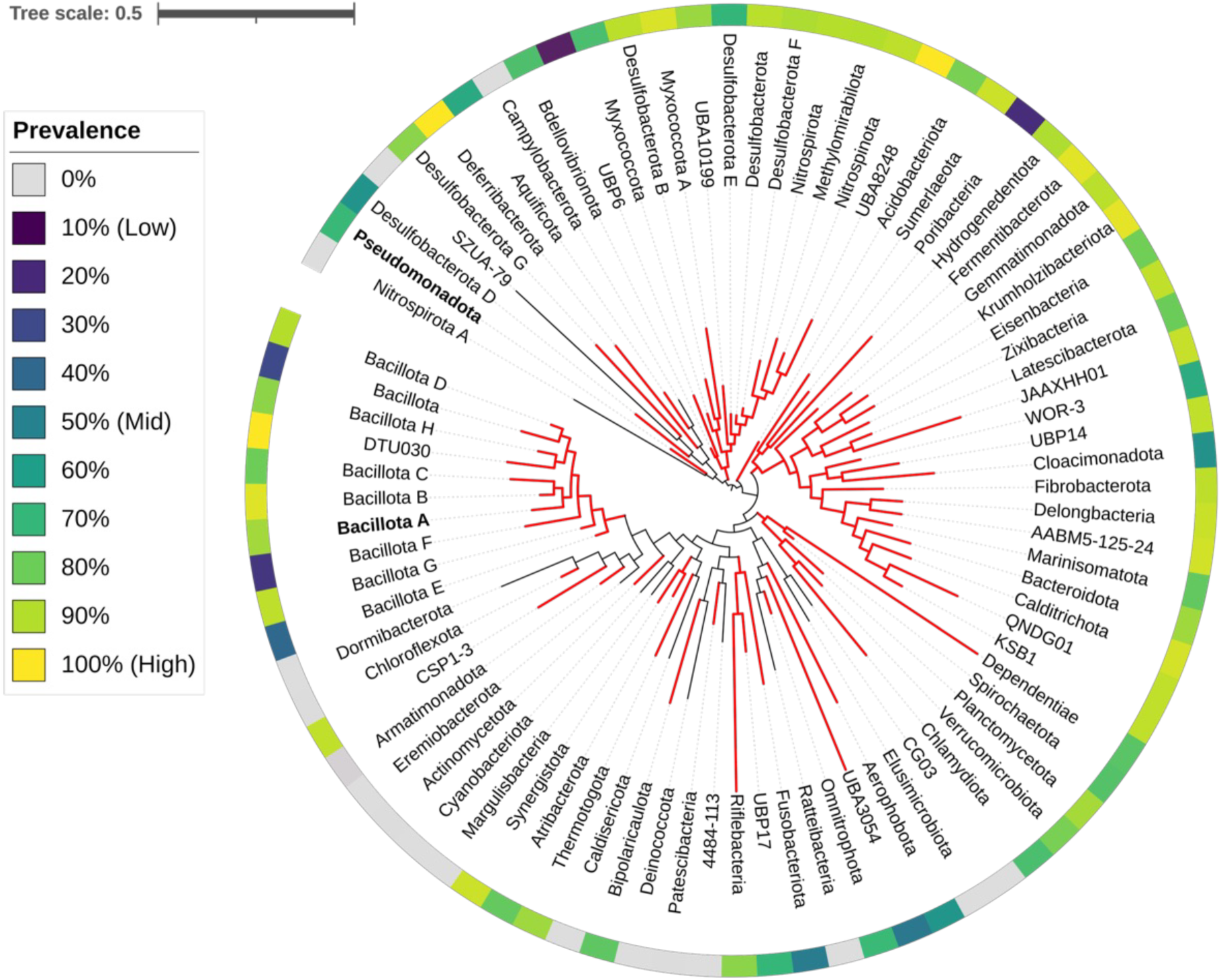
Phylogenetic distribution of YicC across the bacterial domain. The tree represents the evolutionary relationships of major bacterial lineages obtained via AnnoTree and based on the Genome Taxonomy Database (GTDB) Release 214 classifications. The outer heatmap ring indicates the prevalence of YicC homologues, calculated as the percentage of sequenced genomes within each phylum that encode the gene. Red branches highlight clades where YicC is present, whereas black branches indicate lineages where the gene is entirely absent. Phyla containing functionally characterised YicC homologues (Pseudomonadota and Bacillota A) are bold. Among the extensively sampled phyla (>1000 genomes annotated in the database), YicC-like proteins are prevalent across Pseudomonadota (71%), Bacillota (70%), Bacteroidota (87%) and Desulfobacterota (89%), but absent in annotated genomes in Cyanobacteriota, Campylobacterota, and in less than 0.5% of genomes across Chloroflexota, Actinomycetota and Patescibacteria. The scale bar represents 0.5 expected amino acid substitutions per site.

**Appendix Figure S2.**
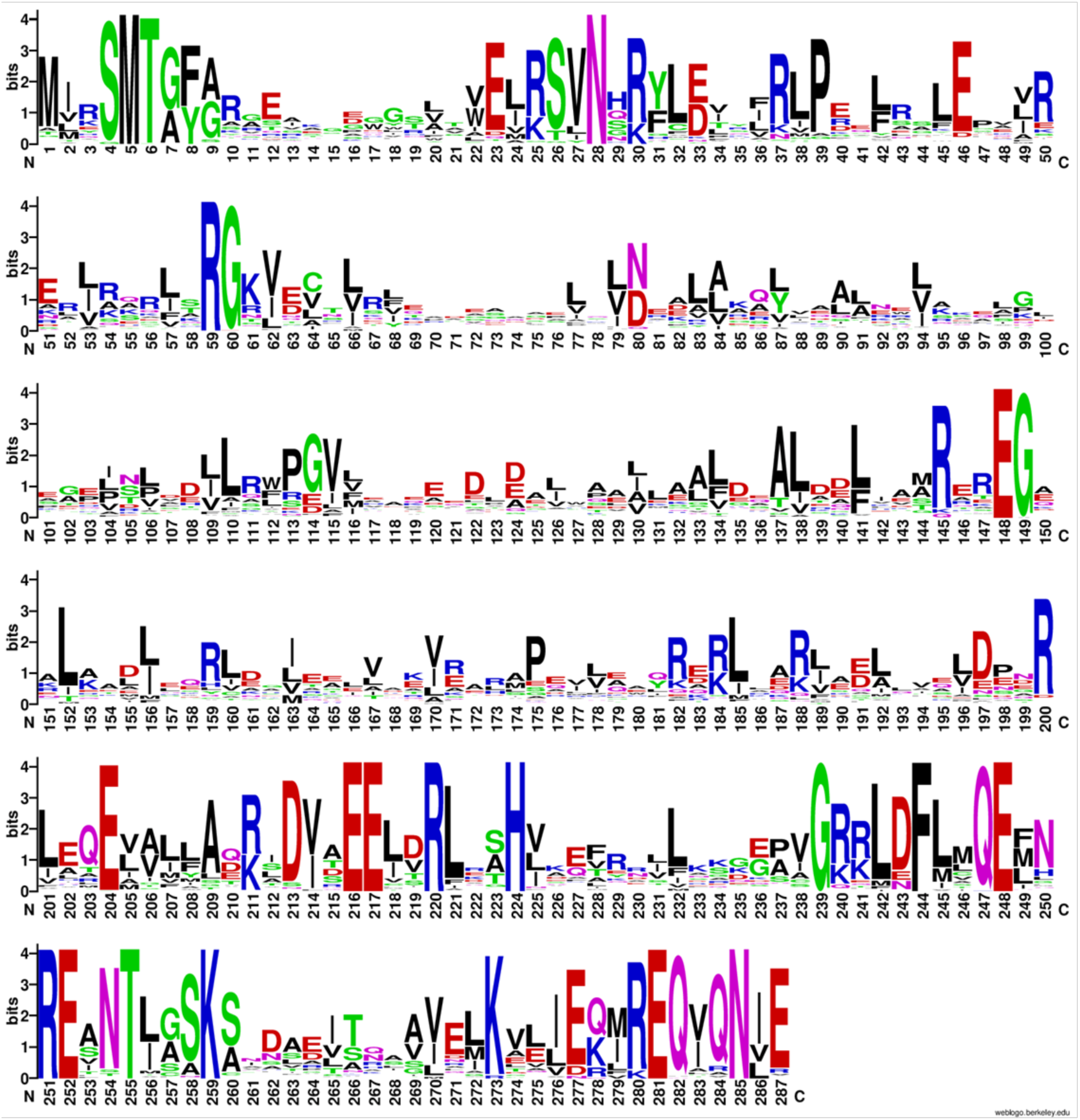
Sequence conservation profile of YicC homologues. Sequence logo generated from a multiple sequence alignment of 87 diverse bacterial YicC homologues using WebLogo. The total height of each amino acid stack indicates the sequence conservation at that position measured in bits, representing the total information content. The relative height of an individual letter within a stack corresponds to the relative frequency of that amino acid at that position. Residues are colour-coded according to their chemical properties: basic (blue: Arg, Lys, His); acidic (red: Asp, Glu); polar (green: Gly, Ser, Thr, Cys, Tyr); neutral/amide (purple: Asn, Gln); and hydrophobic (black: Ala, Val, Leu, Ile, Pro, Met, Phe, Trp). Strict invariance and high conservation across several positions underscore highly conserved structural and catalytic motifs critical to YicC endoribonuclease function across diverse bacterial phyla.

**Appendix Figure S3.**
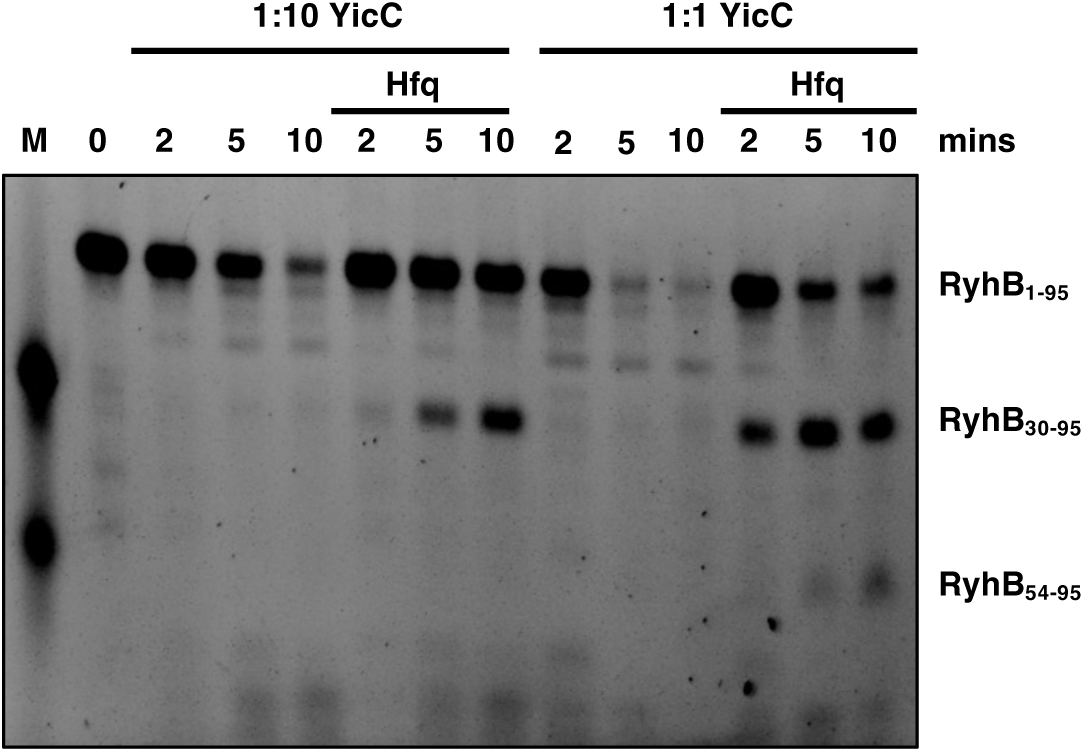
YicC cleavage of RyhB at different enzyme:substrate ratios. *In vitro* degradation assays showing the stability of RyhB sRNA over a 10-minute time course. 200 nM RyhB was incubated with YicC at either 20 nM (1:10 enzyme:substrate ratio) or 200 nM (1:1 enzyme:substrate ratio) in the absence or presence of 200 nM Hfq. *In vitro* transcribed RyhB_30-95_ and RyhB_54-95_ fragments are loaded as standards (lane M).

**Appendix Figure S4.**
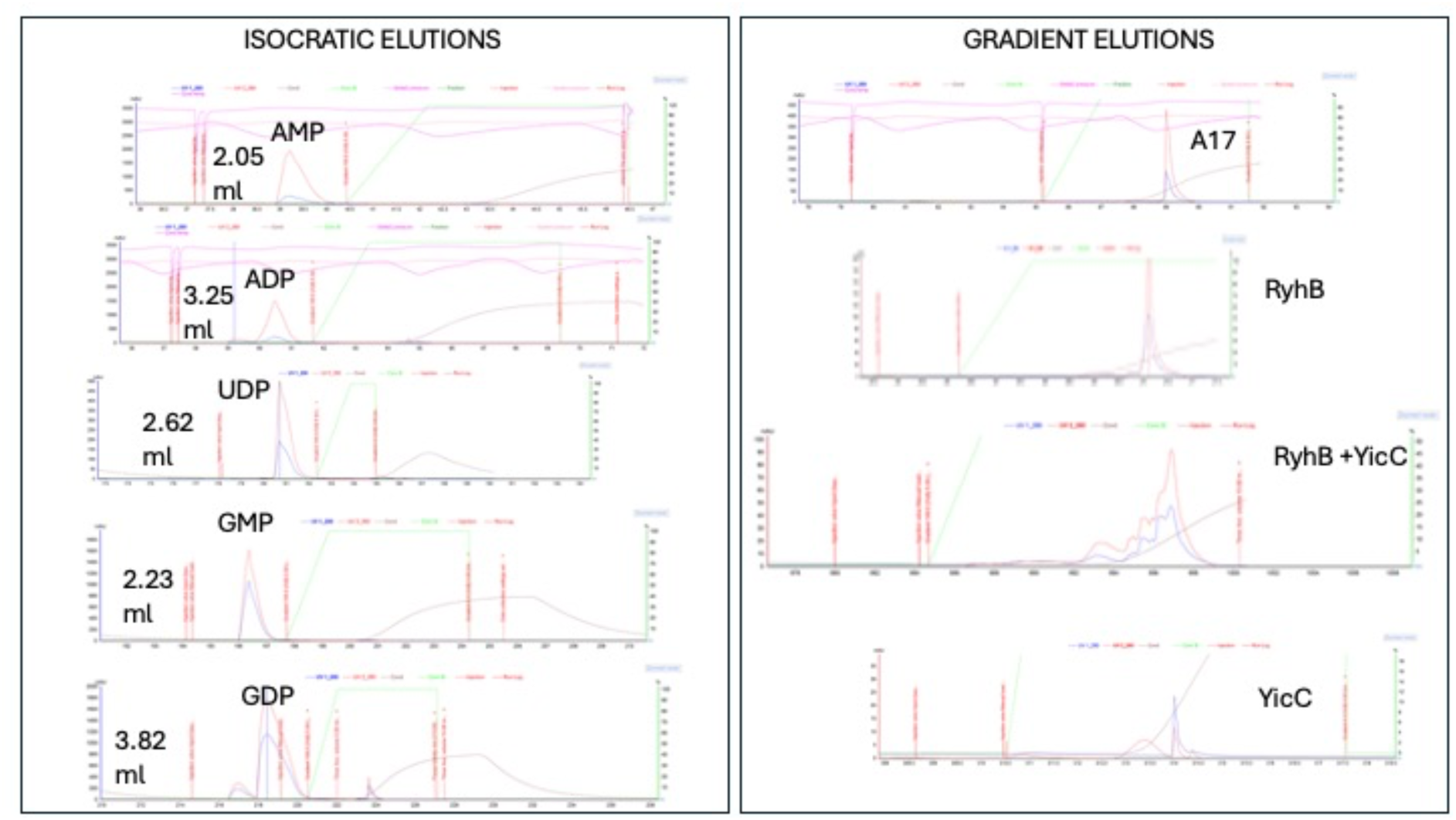
YicC cleavage does not generate nucleoside mono- or diphosphates. Chromatographic profiles of nucleotide mono- and diphosphates (left panel) and RyhB and YicC treated RyhB (right panel). The right panel also shows profiles for A17 and YicC. The chromatograms were preparing using a Dionex DNAPac PA-100 exchange column. The column was equilibrated with 50 mM Tris pH 7.5, 0.5 mM EDTA, and a gradient applied to 1 M NaCl. The nucleoside mono-and diphosphates elute isocratically at the indicated volumes, while the RNAs and digest product isolate in the gradient.

**Appendix Figure S5.**
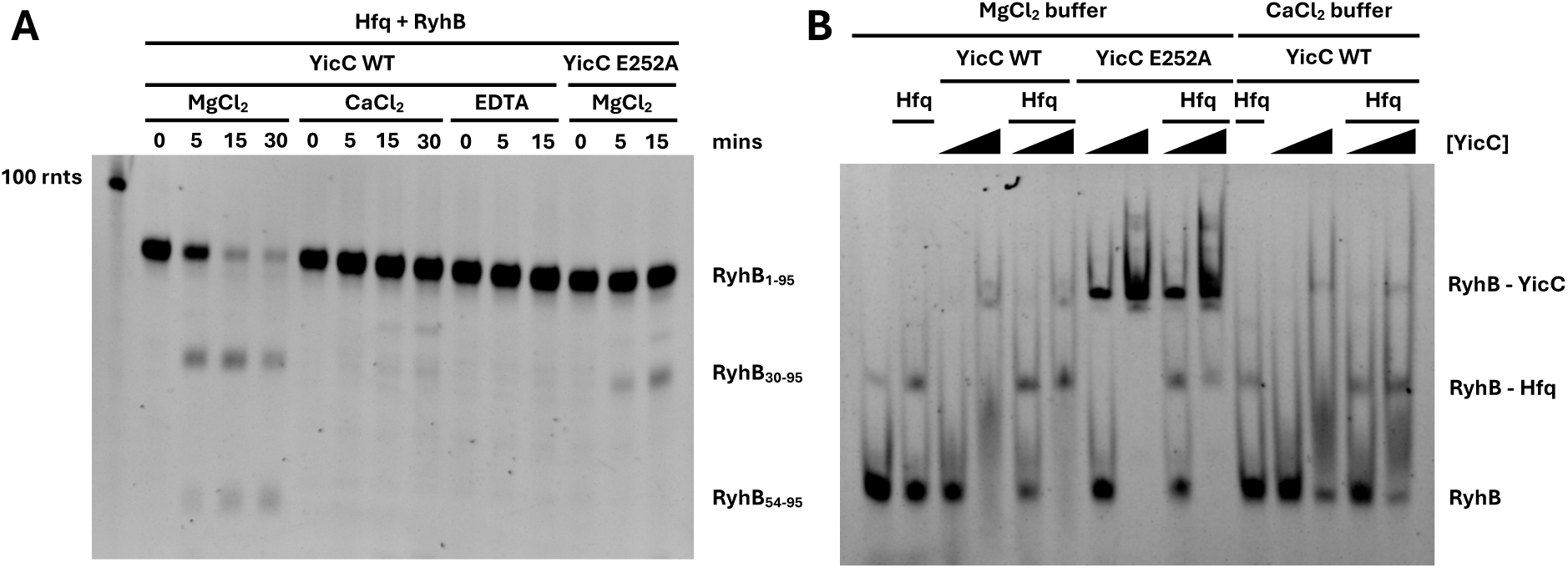
Reconstituting a stable YicC-RyhB complex for structural studies. **(A)** *In vitro* degradation assay showing YicC-mediated cleavage of Hfq-bound RyhB. Wild-type YicC rapidly cleaves RyhB when MgCl₂ is present but not when it is substituted for CaCl₂. A catalytically impaired mutant, YicC E252A, shows significantly reduced cleavage activity, making it suitable for trapping a pre-cleavage state. **(B)** Electrophoretic mobility shift assay (EMSA) assessing YicC-RyhB complex formation. Wild-type YicC cleaves RyhB in MgCl₂ and, thus, does not form a stable interaction. The YicC E252A mutant forms a stable complex with Hfq-RyhB in MgCl₂. The substitution of MgCl_2_ for CaCl_2_ destabilises the interaction. No stable Hfq-RyhB-YicC ternary complex seems to assemble under the conditions tested.

**Appendix Figure S6.**
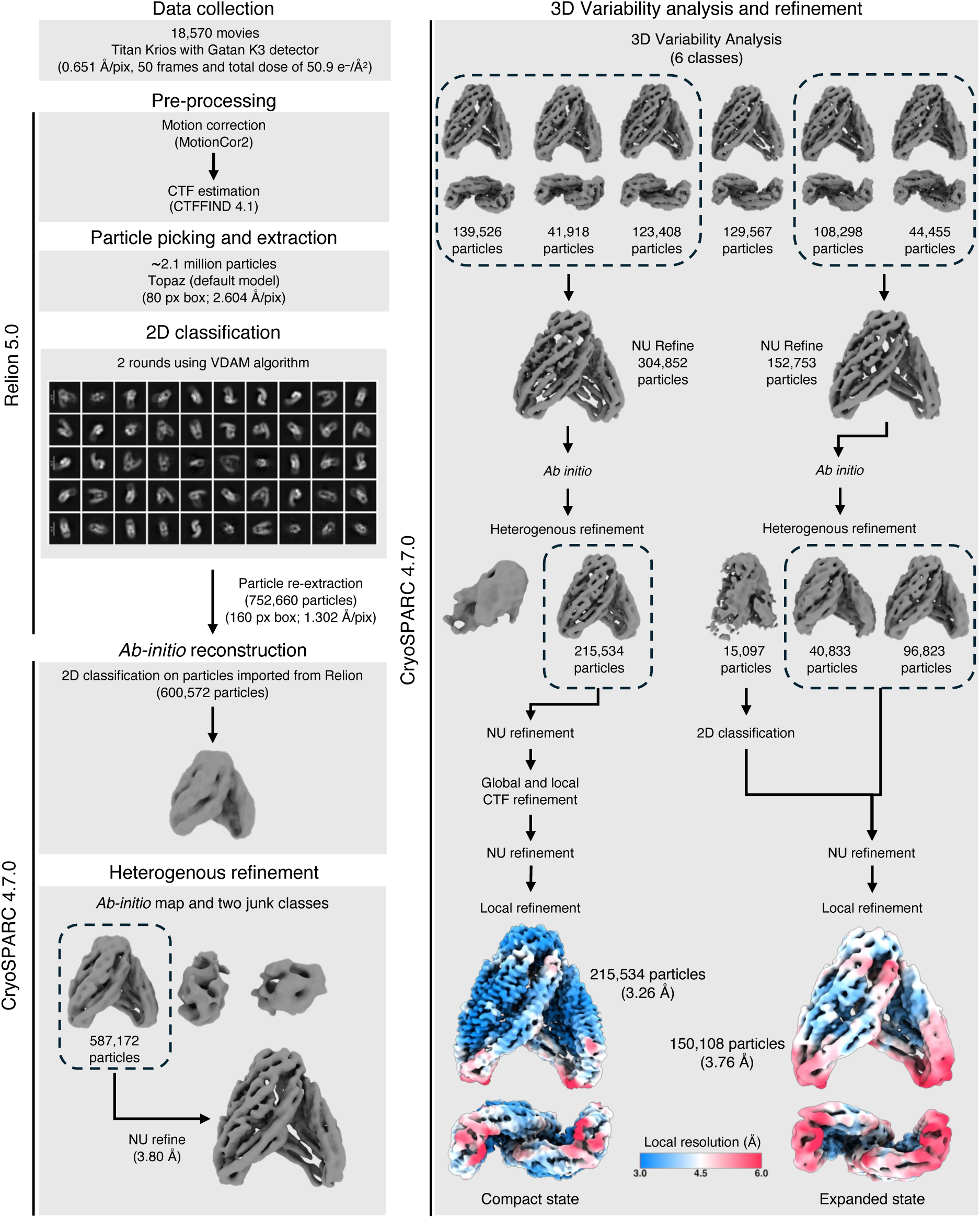
Cryo-EM data processing workflow for apo-YicC. Schematic overview of the image processing pipeline tracking the transition from 18,570 raw micrographs to distinct 3D reconstructions. Pre-processing, Topaz particle extraction, and initial 2D classification were executed in RELION 5.0 and CryoSPARC 4.7.0 to isolate a core stack of 587,172 high-quality particles. To resolve continuous structural heterogeneity, 3D Variability Analysis (3DVA) was used in cluster mode to segregate the dataset into 6 discrete structural classes. Independent Non-Uniform (NU) and local refinement pathways were used to resolve the extreme ends of this conformational landscape: the **Compact State** (215,534 particles; 3.26 Å resolution) and the **Expanded State** (150,108 particles; 3.76 Å resolution). Final unsharpened density maps are displayed at the bottom, coloured by local resolution (3.0 Å to 6.0 Å).

**Appendix Figure S7.**
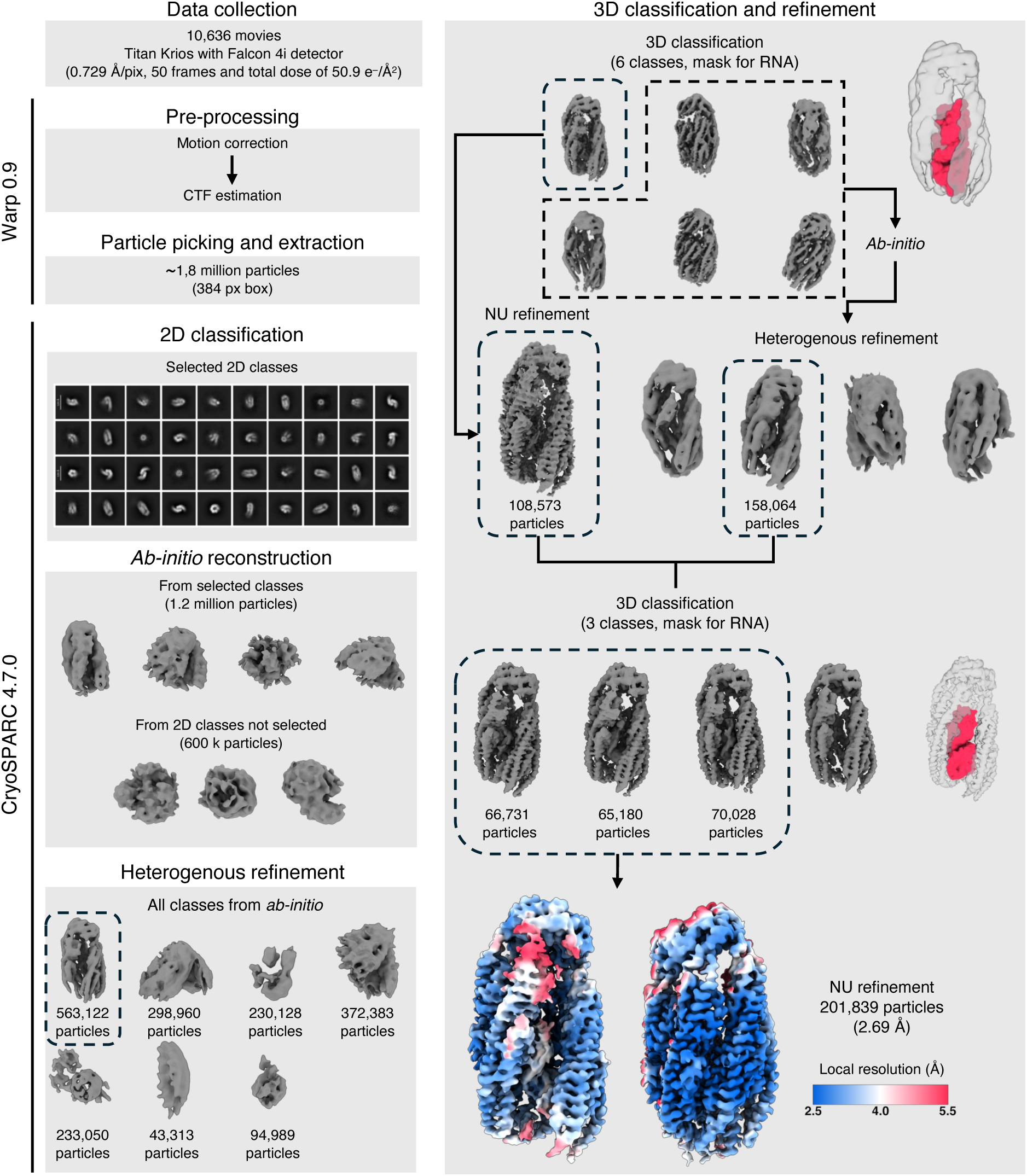
Cryo-EM data processing workflow for YicC-RyhB_1-95_. Schematic overview of the single-particle image processing pipeline tracking the transition from 10,636 raw micrographs to the final high-resolution reconstruction. Pre-processing, CTF estimation, and automated particle picking (∼1.8 million particles) were performed using Warp 0.9. Following initial 2D classification cleanup in CryoSPARC 4.7.0, selected particles (1.2 million) were used for *ab initio* modelling and multi-class heterogeneous refinement to isolate the closed, RNA-bound confirmation (563,122 particles). To isolate high-occupancy complexes, the dataset was subjected to two successive rounds of 3D classification using a focus mask targeting the inner embedding cavity. A final pool of 201,839 particles was selected for Non-Uniform (NU) refinement, yielding a final global reconstruction at 2.69 Å. Final maps are displayed at the bottom, coloured by local resolution (2.5 Å to 5.5 Å).

**Appendix Figure S8.**
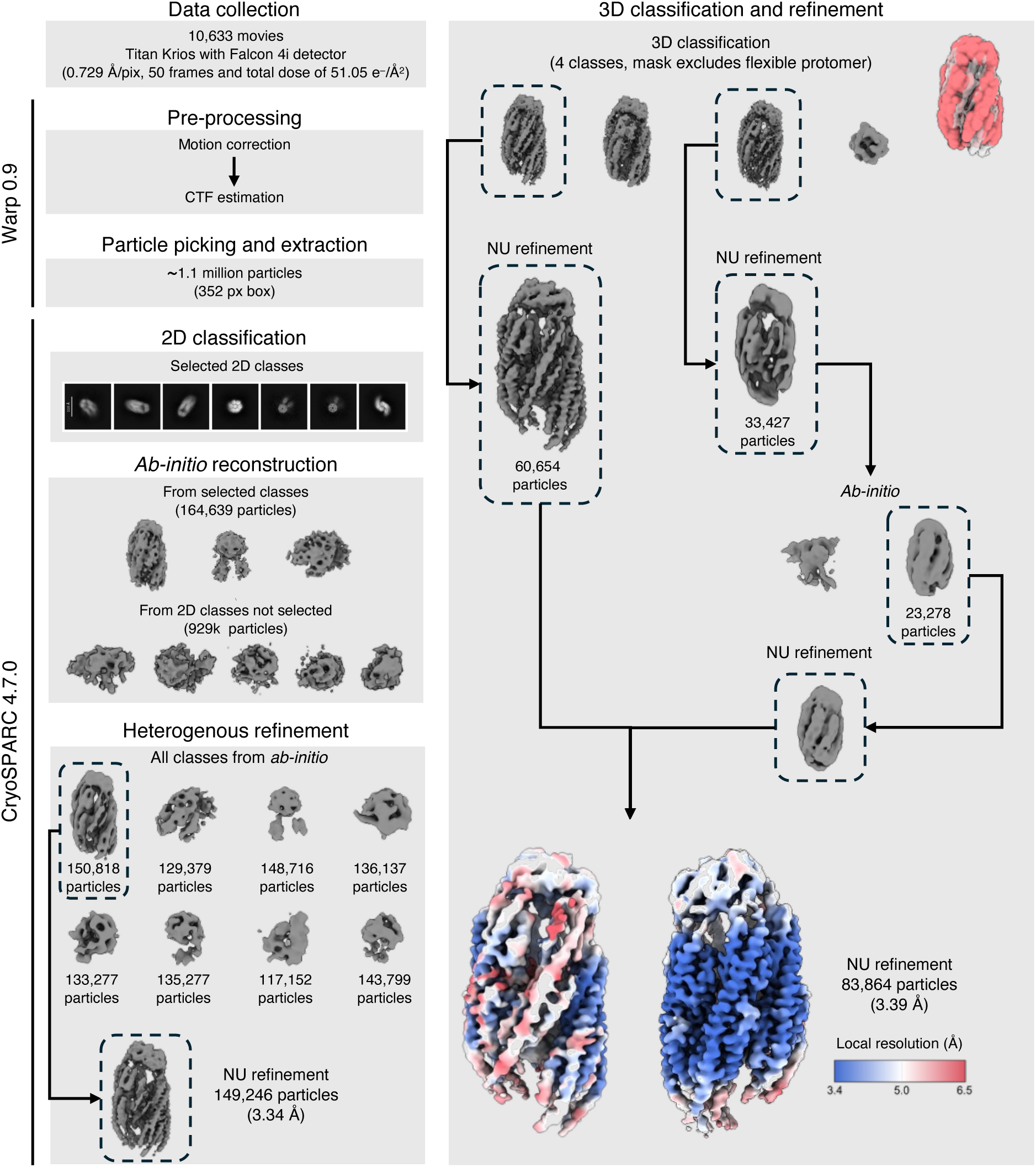
Cryo-EM data processing workflow for YicC-RyhB_19-59_. Schematic overview of the single-particle image processing pipeline tracking the transition from 10,329 raw micrographs to the final high-resolution reconstruction. Pre-processing, CTF estimation, and automated particle picking (∼1.1 million particles) were performed using Warp 0.9. Following initial 2D classification in CryoSPARC 4.7.0, selected particles (164,639) were used for *ab initio* modelling and multi-class heterogeneous refinement to isolate a closed conformation with clear RNA density (150,818 particles). To resolve structural flexibility, the dataset was subjected to focused 3D classification using a mask that explicitly excluded the flexible protomer of YicC. Two distinct classes exhibiting enhanced density for the RNA were selected for further curation and Non-Uniform (NU) refinement, yielding a final core reconstruction of 83,864 particles at 3.39 Å global resolution. Final maps are displayed at the bottom, coloured by local resolution (3.4 Å to 6.5 Å).

**Appendix Table S1.**
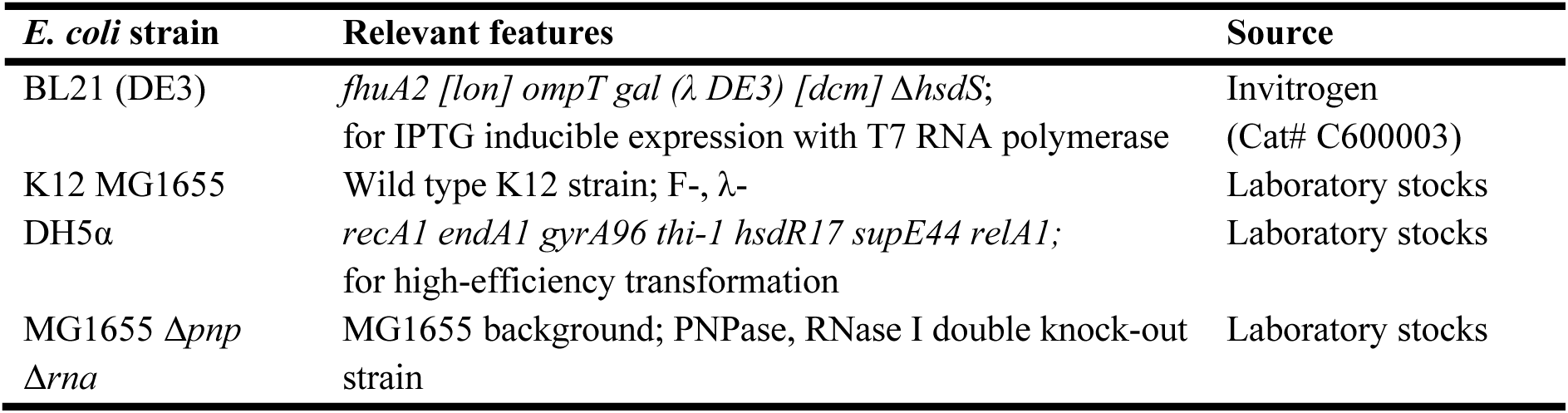
List of *Escherichia coli* strains used in this study.

**Appendix Table S2.**
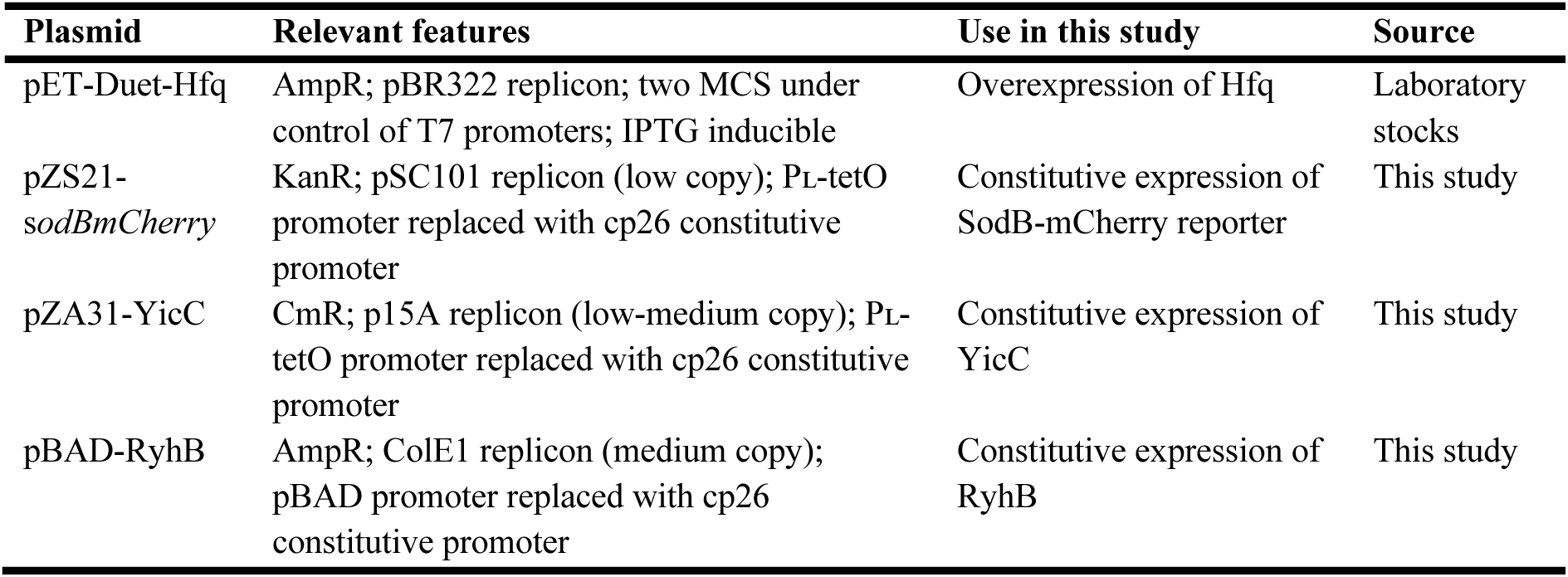
List of plasmids used in this study.

**Appendix Table S3.**
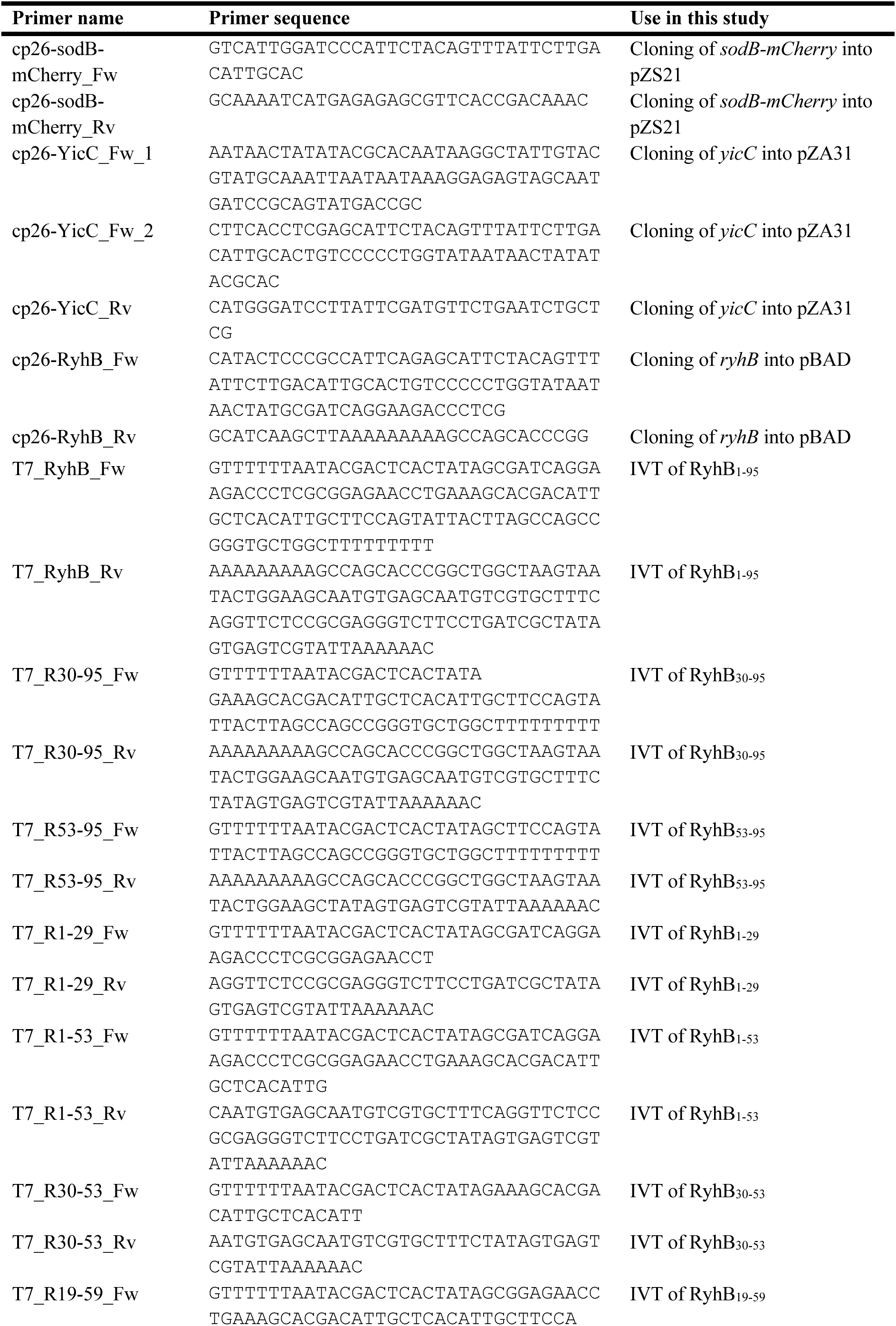
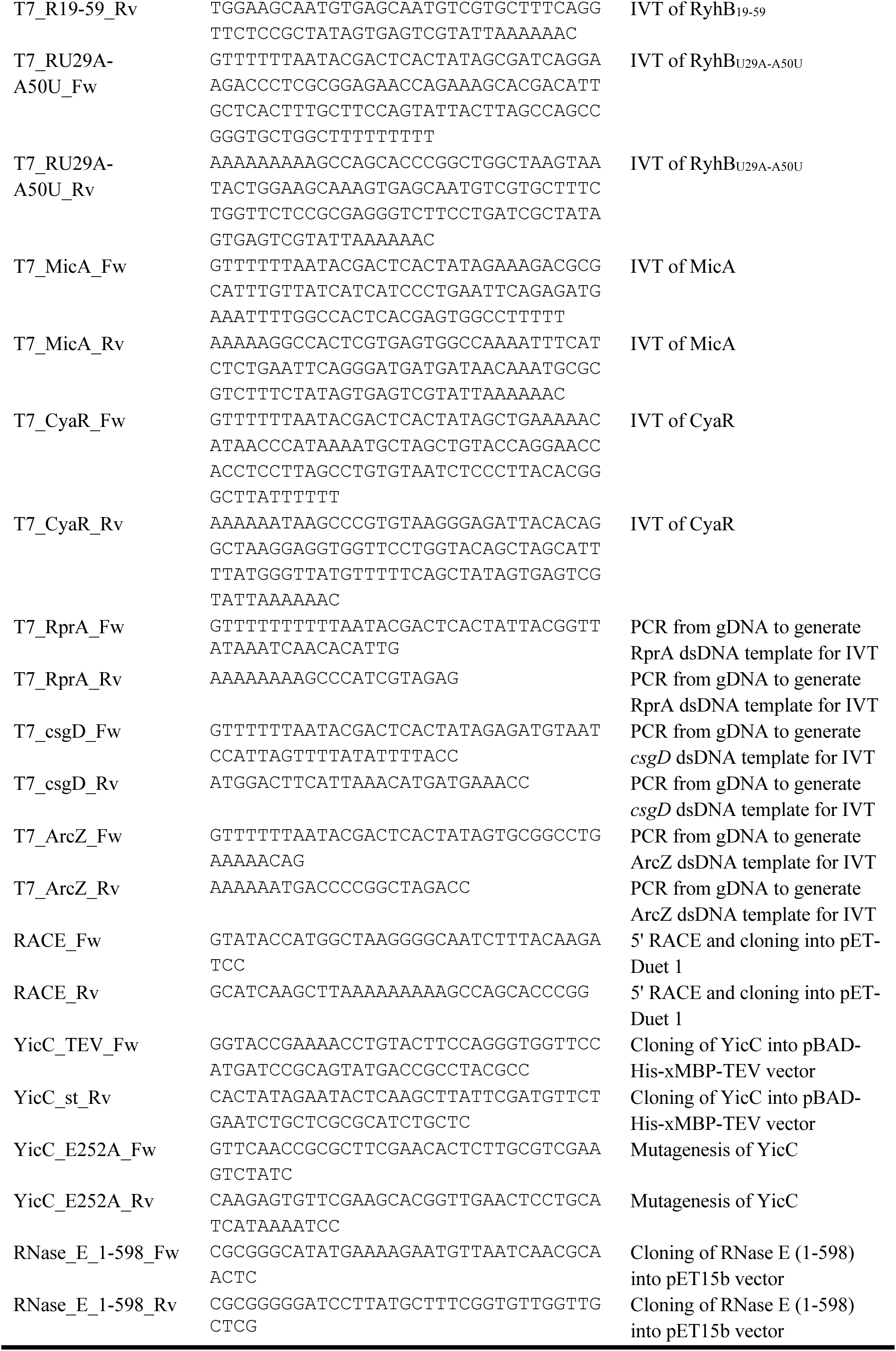
List of primers used in this study. Primers are in 5’ to 3’ orientation.

**Appendix Table S4.** CryoEM data collection and map reconstruction parameters.

| Structure | Apo-YicC |  | YicC-RyhB<sub>1-95</sub> | YicC-RyhB<sub>1-95</sub> |  | YicC-RyhB<sub>19-59</sub> |
| --- | --- | --- | --- | --- | --- | --- |
|  |  |  | Dataset 1 | Dataset 2 |  |  |
| Data collection |  |  |  |  |  |  |
| Microscope | FEI Titan Krios |  | FEI Titan Krios | FEI Titan Krios |  | FEI Titan Krios |
| Voltage (kV) | 300 |  | 300 | 300 |  | 300 |
| Detector | Gatan K3 |  | Falcon 4i | Falcon 4i |  | Falcon 4i |
| Nominal magnification | 130,000x |  | 165,000x | 165,000x |  | 165,000x |
| Pixel size (Å) | 0.651 |  | 0.729 | 0.729 |  | 0.729 |
| Exposure (s) | 1.34 |  | 4.39 | 4.39 |  | 4.39 |
| Dose electrons/pixel/sec | 16.1 |  | 6.66 | 6.18 |  | 6.18 |
| Frames | 50 |  | 30 | 50 |  | 50 |
| Electron dose, total (e<sup>-</sup>/Å<sup>2</sup>) | 50.9 |  | 55.02 | 51.05 |  | 51.05 |
| Electron dose, per frame (e<sup>-</sup>/Å<sup>2</sup>) | 1.02 |  | 1.83 | 1.02 |  | 1.02 |
| Defocus range (µm) | -0.5 to -2.5 |  | -0.6 to -1.8 | -0.6 to -1.8 |  | -0.6 to -1.8 |
| Number of micrographs | 18,570 |  | 9,869 | 10,636 |  | 10,329 |
| Reconstruction | Expanded | Compact | State 1 | State 1 | State 2 | State 1 |
| Particles used | 150,108 | 215,534 | 133,450 | 201,839 | 201,839 | 83,864 |
| Final resolution (Å), FSC<sub>0.143</sub> | 3.76 | 3.26 | 3.08 | 2.69 | 2.69 | 3.39 |
| Map-sharpening B factor (Å<sup>2</sup>) | -164.4 | -99.2 | -92.7 | -70.1 | -70.1 | -80.54 |
| PDB code | 30AB | 29ZL | 30BE | 30EK | 30ET | 30FI |
| EMDB code | 57480 | 57472 | 57534 | 57665 | 57665 | 57694 |

